# Systemic bio-inequity links poverty to biodiversity and induces a conservation paradox

**DOI:** 10.1101/2025.05.13.653456

**Authors:** Conor Waldock, Van Thi Hai Nguyen, Margaret Awuor Owuor, Anna Samwel Manyanza, Ole Seehausen

## Abstract

1. Biodiversity is declining globally while inequity is growing, and poverty rates are not improving. Global sustainable development and conservation initiatives aim to address biodiversity loss and poverty simultaneously.
2. Through text analysis of global biodiversity policies, we identified a consistent narrative that countries with high biodiversity are expected to use this natural capital to reduce poverty. Theoretically then, should higher biodiversity lead to higher prosperity? Clearly countries with higher biodiversity often experience higher poverty rates, so we explore how this paradox emerges using a path analysis integrating multiple mediating factors represented in biological, historical, and socio-economic data.
3. We find that historical colonial exploitation was directed towards biodiversity rich regions, and through socio-economic legacies, continues to influence present-day poverty. At a country-scale, biodiversity is associated with higher poverty rates through increasing the likelihood of historical colonisation. Colonisation drove colonised countries towards natural resource exportation with weakened governance and increased poverty rates. We find no statistical support that country-scale biodiversity levels directly influence poverty, either positively or negatively.
4. Moreover, we find that conservation investments did not align with countries’ biodiversity or their economic capacity to protect nature, indicating a conservation paradox. Our work uncovers that conservation initiatives to protect biodiversity ignore that biodiversity is strongly associated as a historical catalyst for modern global inequalities.
5. Our findings quantitatively support the “systemic bio-inequity” hypothesis proposed here. This hypothesis argues that modern biodiversity-poverty linkages are, at least partly, attributed to historical legacies of unequal exploitation of the natural resources provided by biodiversity. This hypothesis aligns with the theory of ecologically unequal exchange. Our findings highlight the problematics of the narrative that natural wealth, and its conservation, will underpin future economic prosperity of countries. This assumption overlooks systemic, historical and institutional drivers of modern poverty.
6. Recognizing that colonial and neoliberal structures have created inverted poverty-biodiversity relationships fundamentally changes how we should approach conservation. A decolonial approach is needed in conservation to address this legacy, whereby conservationists must not only recognise but challenge and replace damaging narratives that overlook past and present inequalities in access to the benefits of biodiversity.

## Introduction

The abundance and diversity of biological organisms are the foundation of human economic prosperity through ecosystem services (IPBES 2019). Yet, biodiversity is not equally distributed across the globe whereby biological richness tends to peak at low latitudes but prosperity tends to peak at high latitudes (Pontarp *et al*. 2019). In addition, biodiversity is globally declining and this loss undermines ecosystem services (IPBES 2019). Biodiversity is expected to underpin well-being, so the spatial pattern that nature’s richness and people’s prosperity are negatively related is paradoxical and indicative of a spatial decoupling between society and nature. This paradox likely arises from unequal exchange of ecosystem services (i.e., ecologically unequal exchange theory; Givens *et al*. 2019; Hornborg 1998) the unequal exchange of material, economic and labour flows between tele-coupled distant regions (Cotta *et al*. 2022; Liu *et al*. 2013, 2015).

Development and environmental policy programs have responded to the contemporaneous decline in biodiversity (IPBES 2019) and rise in absolute poverty and inequality (Hickel 2018a) by addressing both these global challenges together (Carmenta *et al*. 2025; Fisher *et al*. 2020; UN 2015). The Convention on Biological Diversity (CBD) marked a shift in 1992 from ecology-focused nature conservation to a development and poverty reduction approach (Smallwood *et al*. 2022) which was replicated in the Aichi Biodiversity Targets (COP 2010). The most recent set of global biodiversity targets, the Kunming-Montreal Global Biodiversity Framework in 2022 (KMGBF), recognise that biodiversity conservation can hinder local well-being and instead promote biodiversity considerations in all aspects of poverty alleviation and societal development (COP 2022). One core concept underlying such thinking is that biodiversity - and “nature” more generally - provides benefits that increase natural capital and thereby improve inclusive wealth (Polasky *et al*. 2015; Turner *et al*. 2012). Somewhat contrasting this narrative, the IMF and World Bank development policies argue that evidence for environmental Kuznets curves support the idea that the depletion, not the conservation, of biodiversity has historically improved living standards (Leal & Marques 2022; Marques *et al*. 2019; Rashid Gill *et al*. 2018). These policy differences highlights the rich set of diverging theories, empirical evidence and policy responses that exist around biodiversity management, well-being outcomes and poverty alleviation which highlights the complexity of socio-ecological systems and the often unexpected outcomes from their governance (Mace *et al*. 2018).

Biodiversity contributes to human well-being, with benefits occurring through providing food and agricultural diversity, climate and environmental regulation, disease control, emotional wellbeing, and a richer cultural sense of place and being (IPBES 2022). However, relationships can be complex (Howe *et al*. 2013; Reader *et al*. 2023) and the critical question remains: *who benefits and whose biodiversity underpins such benefits?* In practice, economic tools now dominate the management of people-nature relationships by pricing and market formation of biodiversity (Kashwan *et al*. 2021a). Biodiversity is commodified through tourism, raw material extraction, land rights sales, agricultural and biochemical patents, or is monetised through Payment for Ecosystem Service schemes (Billé *et al*. 2012; Engel *et al*. 2008; Ferraro & Hanauer 2014; Kiwango & Mabele 2022; Koot *et al*. 2024; Muradian *et al*. 2013; Salzman *et al*. 2018). This monetary-nature linkage has recently been solidified through initiatives to mainstream nature in global economic systems, often justified as necessary to conserve biodiversity (Engel *et al*. 2008; IIED & UNEP-WCMC 2017; Milne 2022).

In general, the discourse of UN meetings and policies has increasingly blended language between business and nature, which constructs the ideological framing of any development solutions (Drury et al. 2022), a framing that often hinders transformative change (IPBES 2024). This economic narrative supports CBD policies that biodiversity and prosperity can, and “should” in a normative sense, be positively related (Drury *et al*. 2022; Mace 2014). This framing masks a critical problem and is contested as it arises from Western concepts of nature protection, often marginalises less powerful rural actors and does not address the inequitable global economic structure that produces biodiversity loss and poverty in the first instance (Büscher & Fletcher 2019; Castree 2003; Foster 1999; Foster *et al*. 2010; Holleman 2018; Kashwan *et al*. 2021a; McAfee 1999; Milne 2022). Therefore, to improve global policy agreements, we must first understand how and why poverty and biodiversity are globally interlinked, and whether current conservation approaches are distributed equitably.

The long history of global political and economic power imbalance remains rarely acknowledged in the natural sciences. Yet, this history structures the ecologically unequal exchange which systemically decouples social (herein economic prosperity in the Global North centre of empire) and ecological systems (herein biodiversity in the Global South periphery of empire) (Dorninger *et al*. 2021; Infante-Amate & Krausmann 2019) - generating a “metabolic rift” (Foster et al. 2010) and leading to poor conservation outcomes (Carmenta *et al*. 2023a). These power imbalances now frame global economic and environmental decision making through inter-governmental organisations and the often hegemonic values they represent (Carmenta *et al*. 2025; Drury *et al*. 2022; Nguyen *et al*. 2025). Many economists and historians have argued that 15^th^ to 20^th^ century European colonialism, imperialism and the associated genocidal enslavement of millions of people and other forms of unpaid labour exploitation were foundational to the emergence of this modern global capitalist economic system (Baptist 2014; Berg & Hudson 2023; French 2021; Williams 1944). This system relies on the (often violent) extraction, undervaluation, and modification of natural resources - including the abundance and biodiversity of life - from formerly colonised countries and their transfer to, transformation, marketing and consumption in the colonising countries by means of unequal exchange and trade (Beinart & Hughes 2007; Crosby 2004; Ghosh 2021; Hickel *et al*. 2024; Hornborg 2006; Horne 2017; Infante-Amate & Krausmann 2019). In such a system the benefits of natural and associated labour resource exploitation are moved away from the costs (Wang *et al*. 2024). A well-recognised paradox then emerges: natural resource wealth slows or stalls countries economic development, largely because of exploitative exchange conditions (Acemoglu *et al*. 2001; Rodney 1974; Ross 1999). As such, environmental policies require a strong evidence base to justify the normative position that biodiversity conservation and nature valuation are effective means to reduce poverty (COP 2002, 2022, 2010). Conservation efforts, either direct or indirect, should be first prioritise countries most responsible for global biodiversity loss and the least burden of poverty (Carmenta *et al*. 2023a; Milner-Gulland 2024; O’Brien *et al*. 2024).

Major global biodiversity conservation initiatives, organised through the CBD, view biodiversity-poverty relationships as critical to consider during development. To set the context for our work, we assessed how biodiversity-poverty relationships are viewed in the 2002, 2010 and 2022 COP strategic biodiversity agreements. A narrative exists that links poverty eradication and biodiversity across three major strategic plans arising from the Convention on Biological Diversity (i.e., The Hague 2002, Aichi 2010 and Kunming-Montreal 2022, Table 1, Table S1). To avoid our own possible biases in summarising the sentiment of such relationships, we use large-language models to summarise these three documents with the resulting output stating: “*<u>Protecting biodiversity is essential for poverty reduction, providing sustainable livelihoods,</u> <u>enhancing resilience, and ensuring access to vital ecosystem services. Integrating biodiversity into</u> <u>development strategies can mitigate poverty and foster long-term sustainability</u>*” (see Methods). While these policies represent normative aspirations, they are predicated on an assumed material relationship: that conserving natural capital *can* and *will* generate local wealth. We propose this statistical linguistic representation of the main sentiments of these policy documents can be summarised as the “*biodiversity underpins local wealth*” hypothesis. We test whether the historical and current structural conditions allow this assumed mechanism to function. We thereby test the match between the normative position arising from these global policy documents and empirical country-level data at the global scale.

**Table 1.** Statistical linguistic summary of major COP policy documents related to biodiversity. Note that we used the entire documents in our LLM representations and selected specific to emphasise where such assertions are made in the text directly.

| Document | Example of a direct quote | Linguistic summary of poverty-biodiversity relationships using LLMs |
| --- | --- | --- |
| COP 6 (2002)<br>Strategic Plan for the Convention on Biological Diversity | Section A 3: "In addition to its intrinsic value, biodiversity provides goods and services that underpin sustainable development in many important ways, thus contributing to poverty alleviation" | The Convention on Biological Diversity emphasizes the critical link between biodiversity conservation and poverty eradication. It acknowledges that economic and social development, especially in developing countries, must be prioritized. The CBD highlights the need for financial resources, technology access, and local community support to promote sustainable development and biodiversity conservation. |
| COP (10/2, 2010)<br>The Strategic Plan for Biodiversity 2011-2020 and the Aichi Biodiversity Targets | Annex Section I 3. "[Biological diversity]... contributes to local livelihoods, and economic development, and is essential for the achievement of the Millennium Development Goals, including poverty reduction." | The text emphasizes the interconnectedness of biodiversity and poverty, noting that biodiversity loss disproportionately impacts the poor, who rely on ecosystem services for survival. It highlights that protecting biodiversity contributes to poverty reduction through better health, food security, and livelihoods, urging integration of biodiversity into development strategies. |
| COP (15/4, 2022)<br>Kunming-Montreal Global Biodiversity Framework | Target 14: "Ensure the full integration of biodiversity and its multiple values into... poverty eradication strategies." | The text highlights implicit connections between biodiversity and poverty, emphasizing that biodiversity conservation can support sustainable livelihoods, ensure access to resources, and promote equitable development. It suggests that the Kunming-Montreal Global Biodiversity Framework can help alleviate poverty by fostering sustainable use of biodiversity, improving access for vulnerable communities, and promoting gender equality. |

The current global economic hegemony, rooted in historical processes, described above, makes the benefits of biodiversity continue to systemically accumulate away from countries with attractive resources and is currently a main active driver of poverty (Andreoli *et al*. 2023; Carmenta *et al*. 2023a; Hickel *et al*. 2022a, 2024; Hornborg 2006; Infante-Amate & Krausmann 2019). This could also be framed as a wealth transfer and benefit of externalising environmental costs (Liu *et al*. 2018b; Wiedmann & Lenzen 2018). A transfer of the benefits of nature’s biodiversity and productivity occurs to the Global North (i.e., the core), if the Global South’s (i.e., the periphery) resources are unequally exchanged but the costs of ecosystem service loss is borne in the Global South, thereby increasing poverty (Hickel *et al*. 2022a). To the extent that this hypothesis applies, it predicts a positive association between poverty and biodiversity arising indirectly through colonialism and its systemic impacts on modern economies. We refer to the imbalanced economic benefits of biological diversity - a structural result of historical and contemporary processes - as the “*systemic bio-inequity*” hypothesis.

Here, we first present a correlative analysis of the current relationship between multiple dimensions of a countries’ biological diversity and indices of poverty, finding them to exhibit a well-established positive correlation (Figure 2 and 3). Our major contribution is to test why this global pattern emerges, by using structural equation modelling with 14 biological, social, historical, economic and geographical factors and their interlinkages (Figure 4 and 5). We test whether the two non-mutually exclusive hypotheses explain the relationship between biodiversity and poverty (e.g., the “biodiversity underpins local wealth” and/or the “systemic bio-inequity” hypothesis). Next, we develop a classification of countries based on jointly considering biodiversity, poverty and conservation investment through protected areas (Figure 6). This helps conceptualise responsibilities and inequalities amongst countries’ contributions to global biodiversity targets such as targets 2 and 3 of the KMGBF (2022), target 11 of the Aichi Biodiversity Targets (2011) and Sustainable Development Goals 14 and 15. To reflect the global and national policy development, we focus on country-level properties, while acknowledging that linkages between poverty, biodiversity and conservation can be highly scale- and context-dependent (Zhang *et al*. 2023).

**Figure 1.**
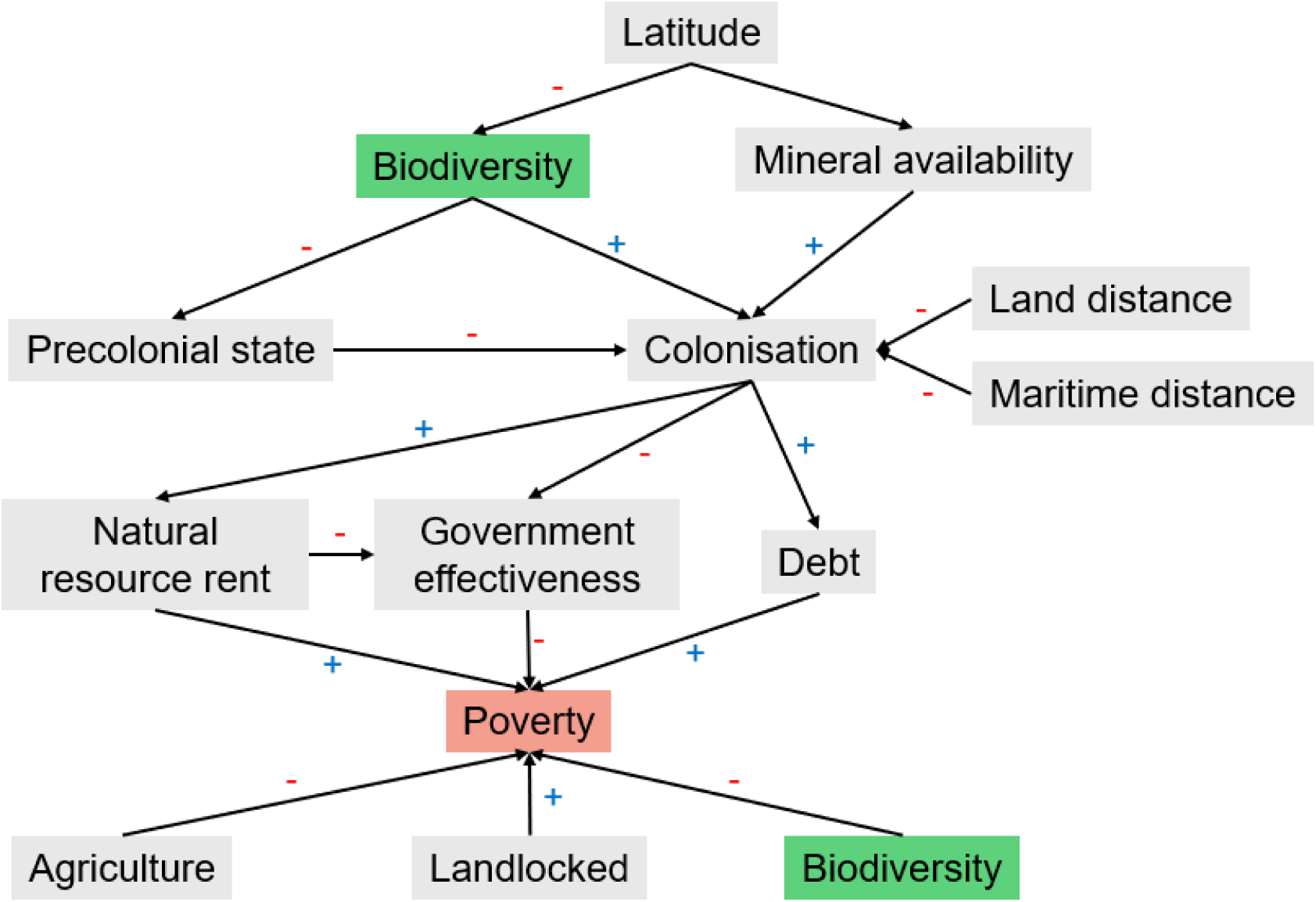
Overview of the hypothesised links tested with data in our structural equation modelling. The direct link between biodiversity and poverty indicates the “biodiversity underpins local wealth” hypothesis, whereas the indirect links between biodiversity and poverty test the “systemic bio-inequity” hypothesis. Note that these hypotheses are not mutually exclusive. Biodiversity related variables are highlighted in green, and poverty variables are in red, to emphasise their placement in this network of socio-economic, geographical, and historical pathways.

**Figure 2.**
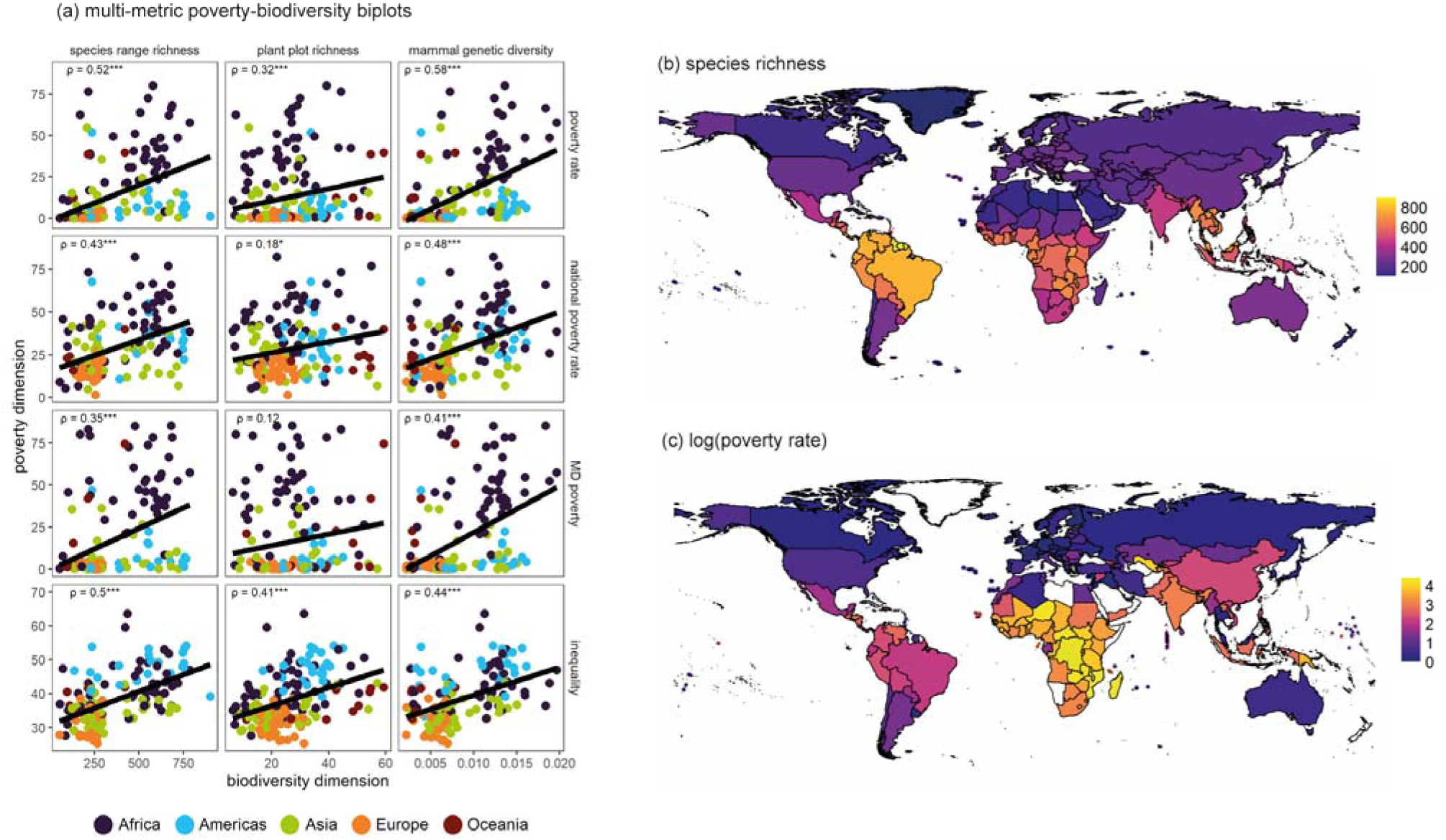
The positive relationship between biodiversity and poverty across multiple metrics (a-c). (a) Shows biplots of biodiversity and poverty metrics with points indicating countries (n=168). Spearman’s rank correlations are indicated in top left with significance of correlation indicated with asterisks (* = p < 0.05, ** = p < 0.01, *** = p < 0.001). (b and c) illustrate the global pattern of biodiversity and poverty with (b) showing terrestrial vertebrate richnes and (c) showing poverty headcount rate using the international poverty line of $2.15 per day.

**Figure 3.**
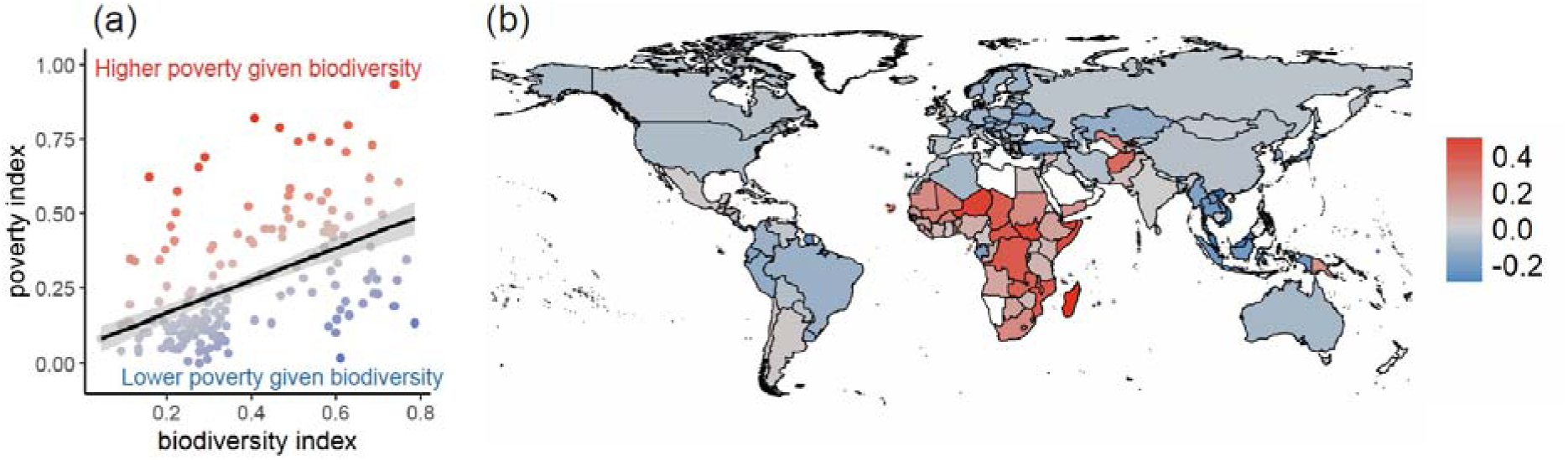
Hotspots of where poverty deviates from the (unexpected) positive relationship with biodiversity. If assuming biodiversity contributes to well-being, then regions in red highlight areas affected by greatest bio-inequity. (a) Shows the relationship between aggregated biodiversity and poverty indicators from Figure 1. (b) Maps the residual value between aggregated metrics of poverty and biodiversity, as shown in (a). The indices were first standardised (0-1) across all countries and then averaged across poverty metrics and biodiversity metrics within each country. Areas and points in blue (poverty < biodiversity) to red (poverty > biodiversity) indicate the residuals of a linear model between these aggregated metrics for poverty as predicted by biodiversity. 95% confidence intervals of the regression coefficient are shown in grey shaded area of panel a.

**Figure 4.**
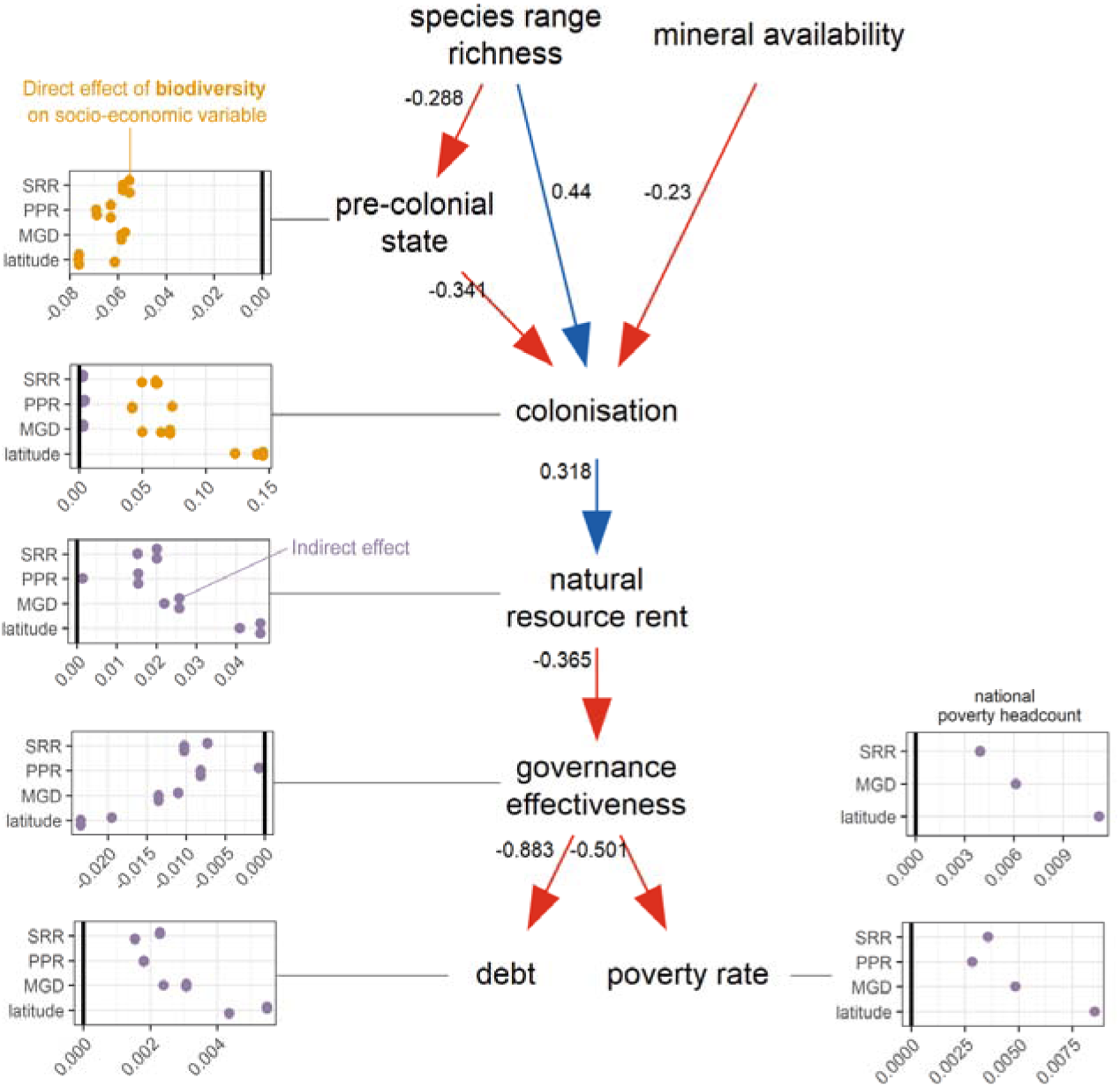
Biodiversity and poverty relationships act within a global socio-economic structure giving rise to systemic bio-inequity, revealed through structural equation modelling. Relationships are presented from our backwards selection of a minimal model based on the full model (Figure 1; Figure S2). Insets show the effects of biodiversity variables (n=3) and sign-inverted latitude on socio-economic metrics. Direct (orange) and indirect (purple) path coefficients were estimated with 1,000 bootstrap samples. Only significant path coefficients are shown, such that missing variable by response combinations are not show in insets. Numbers on edges indicate standardized effect sizes. All path coefficients and indirect effects are significant at p < 0.05. R^2^ values were as follows: pre-colonial state = 0.13, colonisation = 0.42, natural resource rent = 0.10, governance strength = 0.39, debt = 0.75 and poverty rate = 0.4. Biodiversity variables are indicated as SRR = species range richness, PPR = plant plot richness, MGD = mammal genetic diversity. This model structure is statistically supported based on Fisher’s C of 45.49, p=0.18, d.f. = 38.

**Figure 5.**
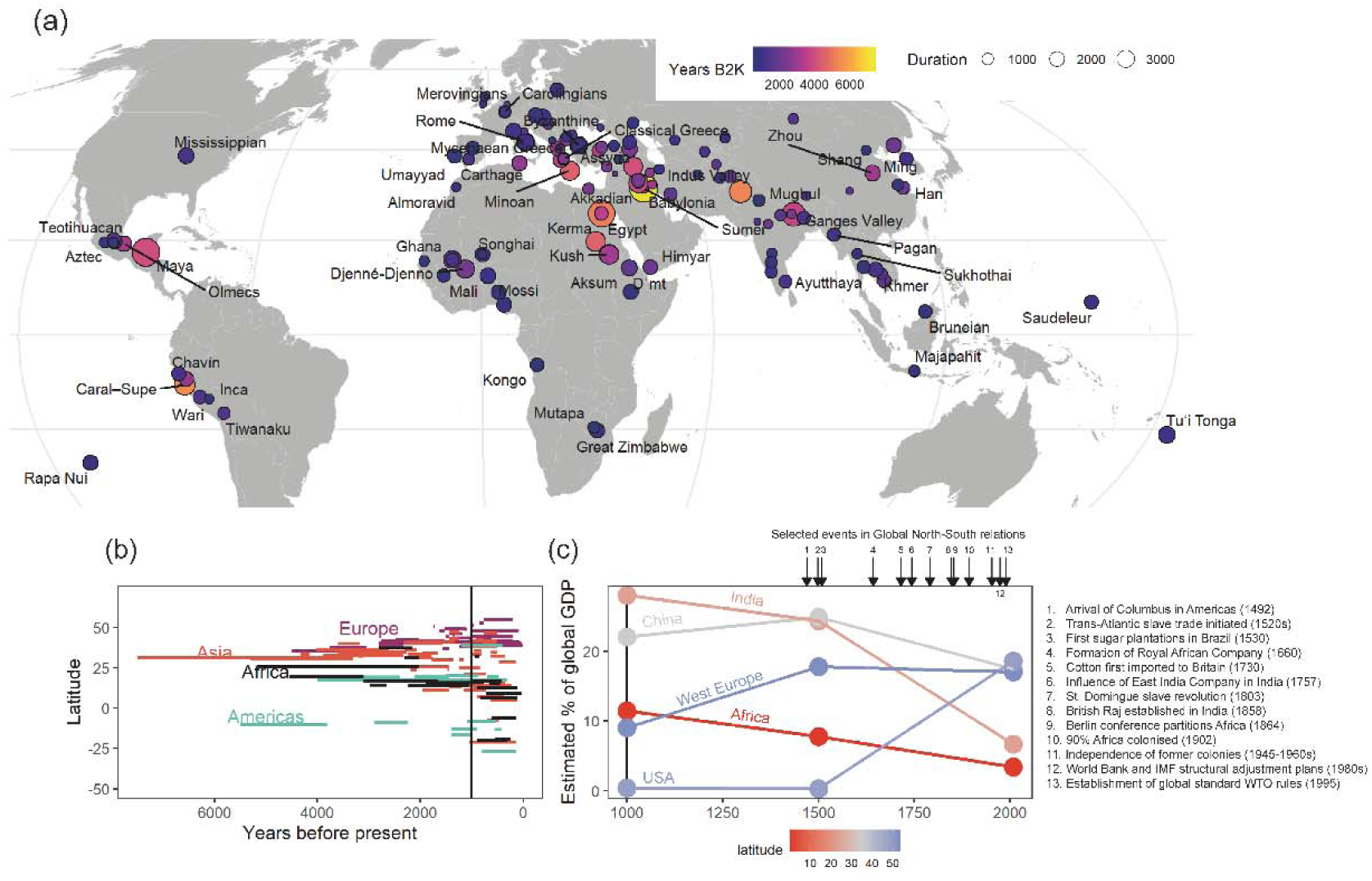
The distribution, emergence and wealth of early complex polities. The global distribution of wealth and earlier emergence of complex polities occurred in regions which now have low proportions of global GDP, these regions were the origins of complex civilisation and had far greater historic GDP. (a) shows the spatial distribution of civilisations exhibiting large infrastructure, cities and complex political systems. The colour of points indicates the years before present when the polities are estimated to have formed, in years before 2000 CE, and the size indicates the duration of the polity. (b) The emergence latitude and time of the polities with colours indicating the geographic regions. The horizontal line is at year 1000, where data on GDP across 5 countries and regions is available for comparison in panel (c). Panel (c) indicates the change in proportional global GDP from year 1000 CE to 2008 CE based on (Maddison 2007). We also indicate major historical events that coupled North-South ecological and economic relationships. Colour in (c) indicates the mean latitude of the region or country.

**Figure 6.**
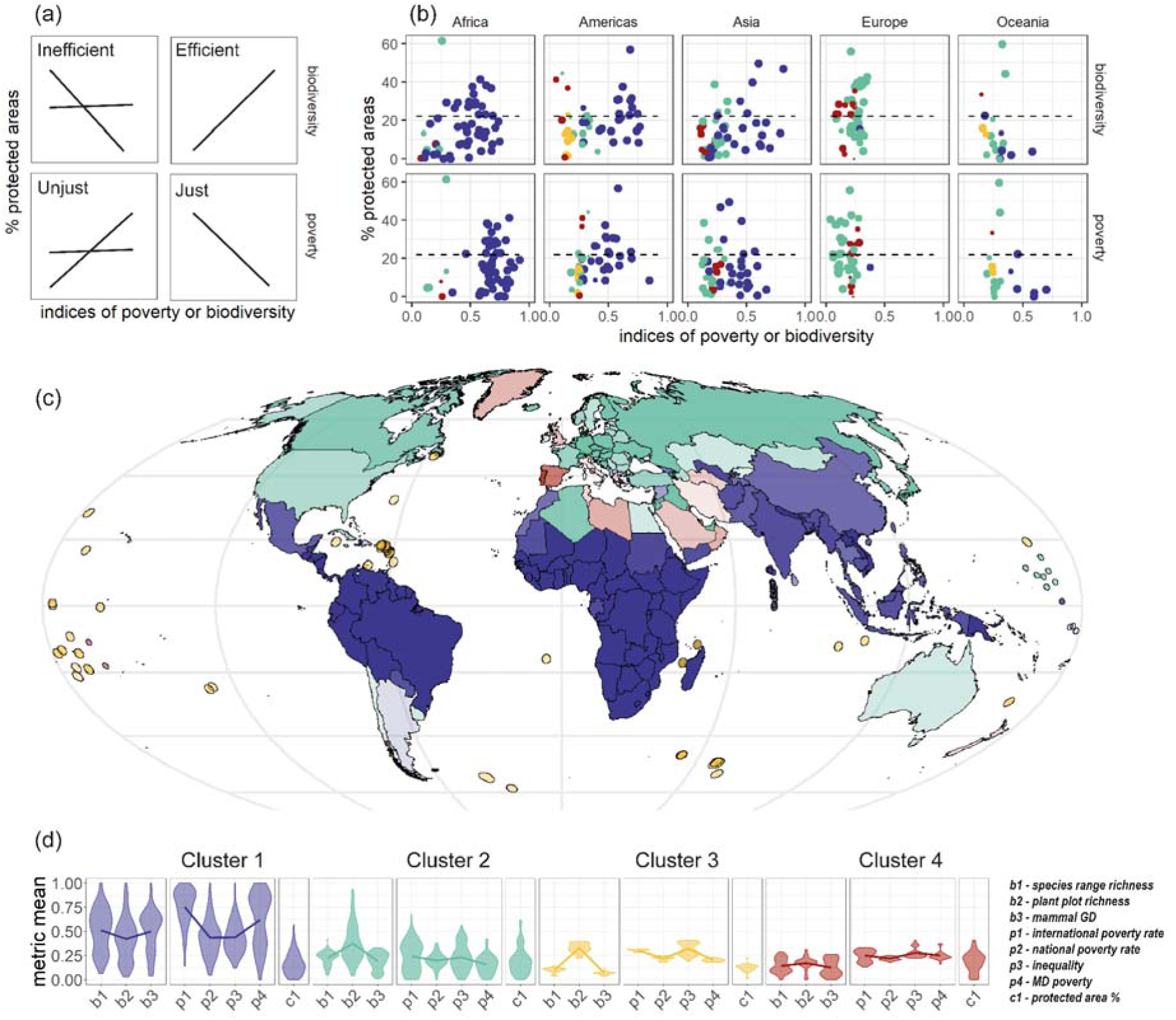
The relationship between conservation investment, biodiversity and poverty across five continental regions. (a) The expected marginal relationships between biodiversity, poverty and protected areas. We highlight whether the global conservation system is efficient and just based on these relationships. (b) Empirical relationships between biodiversity and poverty indices with % protected area coverage for each country, indicated by points. Generalized linear models found no support for significant relationships in any of these regions, with an intercept only model the best supported (dashed lines). Colours indicate the cluster identity of countries based on unsupervised learning using Gaussian mixture models using all metrics of biodiversity, poverty and conservation. (c) Shows a global map of each cluster identity. Note that group 3 is exclusively island countries, characterized by very low species and genetic richness of vertebrate animals. For easier visualisation, we buffered these points by 100km. (d) Indicates the mean values and density distributions of the 8 metrics of biodiversity, poverty and conservation which were used as quantitative inputs to the GMM for the 4 country clusters.

## Methods and materials

### Large language sentiment assessment of United Nations Conference of Parties Convention of Biological Diversity policy documents

We aimed to produce a linguistic statistical representation of the relationship between poverty and biodiversity from major policy documents in biodiversity science. We obtained PDF versions of the Rio 1992, Aichi 2010 and Kunming-Montreal 2022 documents and used *pdftools* (version 3.5.0 (Ooms 2025)) in R to extract all text from these documents. For each document separately, we used the open-source mistral-nemo (24.07) Large Language Model (LLM) to analyse these documents. We tested a range of temperature parameters (0.1, 0.2, 0.3) and token sizes (1,000 and 10,000). We assigned the LLM the following role: “*You answer the following question only based on the provided text, what is the relationship between biodiversity and poverty?”.* For each temperature and token size combination, we provided the text of each policy document as a message and saved the returned output summary. We tested whether the LLM used its “role” to provide the returned message by inputting two dummy text objects: i) the poem “*Because I could not stop for Death*” by Emily Dickenson and ii) a series of 20 random sentences of 100 words using *OpenRepGrid* package (version 0.1.17 (Heckmann 2024)). Analysing both these dummy summaries returned statements confirming these documents were not related to biodiversity or poverty. We saved all output summaries of the provided documents across all parameters and provide a selected set of these in Table S1. For each policy document, we combined all of text summaries from mistral-nemo across all parameter settings and obtained a meta-summary of the document summaries using chatGPT-4.0 turbo, for convenience, requesting the following “*Summarise the following text in 50 words, emphasising the relationship between poverty and biodiversity:*”. We also input all mistral-nemo summaries as a single text string into chatGPT-4.0 for a single summary provided in the text of the main manuscript.

### Data synthesis

We compiled multiple global scale biodiversity datasets representing intraspecific genetic to community scales of biological diversity covering multiple taxonomic groups. We extracted country mean values of: i) terrestrial species richness for amphibians, birds, mammals and reptiles by summing overlapping range polygons to a global harmonised raster layer (IUCN Red List Data 2022-2); ii) terrestrial mammal genetic diversity of co1 and cytb mitochondrial genes comparing the mean number of nucleotide differences per site within a species for >2,000 species and >54,000 sequences (Theodoridis *et al*. 2020); and iii) plant local scale species richness based on >400,000 vegetation plots in a small-scale delimited area ((Sabatini *et al*. 2022); w3_tile_joint_sr1000.tif). Note that the latter two global layers are generated by modelling environmental correlates of diversity and projecting these spatially to provide a globally continuous layer (see original publications for details). Our approach aimed to broadly indicate global-scale spatial patterns in biodiversity using measures of biodiversity that are comparable between countries. We acknowledge that substantial biodiversity knowledge shortfalls exist in many countries but assume that global scale gradients are well represented herein (Hortal *et al*. 2015). We intentionally did not aim to investigate within-country biodiversity variation where locally collected data would be required. We also note that the high dimensionality of biodiversity prevents any attempt at being comprehensive, we aimed instead to be find indicative biodiversity metrics.

We compiled four metrics of poverty and inequality from the World Bank Group’s data portal: poverty headcount based on an international poverty line; poverty headcount based on national poverty lines; multidimensional poverty index; and, income inequality using the Gini index. Poverty headcounts indicate the % of the country living below the international poverty line of $2.15/day based on 2017 Purchasing Power Parity. Poverty headcount using national poverty lines use country-specific national poverty lines provided by household surveys by each country’s government sources or World Bank staff which reflect cost of living differences between countries. Some economists argue that international poverty lines inflate estimates of poverty reductions in recent decades compared with national poverty lines, so we provided here both for consideration (Hickel 2018a; Pogge & Reddy 2006; Ravallion 2010). Poverty measured in monetary terms does not represent non-monetary forms of poverty (Alkire *et al*. 2015) so we also used the multidimensional poverty index that equally weighs monetary poverty, school enrolment rates, primary education completion, access to clean drinking water, access to sanitation and household electricity supply. Using this combined poverty index, individuals are considered poor if in the lowest ⅓ of the combined weighted index, and the country metric is expressed as the % of individuals indicated as poor. Finally, economic inequality is measured using the Gini index to calculate the wealth share amongst individuals within the country: where 0 is perfect equality in wealth between individuals and 100 is complete inequality. For each metric and country, we averaged all available poverty metrics from the year 2000 to the most recent available date. Poverty headcounts and Gini inequality were available from the World Bank Poverty and Inequality Platform using the R package *pipr* ((Fujs *et al*. 2023); version 1.0.0) and we obtained the multidimensional poverty index from the Multidimensional Poverty Measure database (8th edition, circa 2021) available at https://www.worldbank.org/en/topic/poverty/brief/multidimensional-poverty-measure which are based on the World Bank’s Global Monitoring Database.

We focused on protected areas as an indicator of conservation investment. These are defined by the International Union for Conservation of Nature as “a clearly defined geographical space, recognized, dedicated and managed, through legal or other effective means, to achieve the long-term conservation of nature with associated ecosystem services and cultural values”. A key policy in the KMGBF is furthering protected area coverage to 30% (COP 2022), and protected areas are a key focus of multiple NGO’s and intergovernmental organisations (IUCN and United Nations Environmental Program World Conservation Monitoring Centre attempt to compile global data to monitor protected area coverage and quality). For convenience, we used the World Bank’s DataBank to obtain country level calculations from the Protected Planet website (https://www.protectedplanet.net/) of % of total land area protected where protected areas are of at least 1,000 hectares (for full methods see https://databank.worldbank.org/metadataglossary/world-development-indicators/series/ER.LND.PTLD.ZS). For some countries there were multiple years available for protected area %, and we took the value of the most recent year available.

We compiled the above data into a country-scale dataset with a total of 168 countries with no missing data for biodiversity metrics and poverty metrics. To test for general patterns of association between country-level poverty and biodiversity, we perform a Spearman’s rank correlation between each pairwise comparison of metrics, giving 12 comparisons (3 biodiversity metrics x 4 poverty metrics) and report the correlation statistic (ρ) and p-value. Spearman’s rank correlation is robust to data being on different scales, having non-linear relationships, and having non-normal distributions. We also calculated a linear model of poverty metrics as a response variable and biodiversity metrics as an explanatory variable, from which we calculate the coefficient and r^2^ statistic. We next calculated an index for each of the metric groups for poverty and biodiversity, by combining all metrics within a group. To do so, we rescaled all metrics between 0-1 and calculated the mean of the rescaled metrics within poverty and biodiversity metric groupings. Next, we calculated the residuals of a linear model between the poverty index and biodiversity index, this residual indicates whether a country’s poverty index is higher or lower than expected given its biodiversity index and the global relationship. Here, we aim to only summarise the broad relationship between variables and do not account for mediating factors and their influence on the effect of variables, which is addressed in the next section.

### Structural equation model of poverty-biodiversity relationships accounting for socio-economic history

We tested how biodiversity is related to poverty using a structural equation model that comprised of multiple modern and historical socio-economic and environmental factors. We used data from (Ertan *et al*. 2016) which provides an overview of the quantitative links between historical factors and the emergence and timing of colonisation. From this publication we obtained country-level data on colonisation, precolonial state formation, latitude, maritime and land travel distance, and landlocked status. Ertan *et al*. consider a country as colonised if > 20% land area was controlled by Belgium, England, France, Germany, Italy, Netherlands, Portugal or Spain between 1462 and 1945 (see Appendix of Ertan *et al*. for full descriptions and justifications). Differing from Ertan *et al*. we considered settler colonies as not colonised, as our aim is to explain modern day poverty whereas settler colonies institutionally benefited from colonisation in terms of economic wealth (Acemoglu *et al*. 2001). We follow Ertan et al in considering Armenia, Azerbaijan, Georgia, Kazakhstan, Kyrgyzstan, Turkmenistan, Uzbekistan, Lebanon, Syria, Isreal, Jordan and Iraq as non-colonised as these were colonised very late, in addition to Ethiopia which was colonised late and for a brief period. Precolonial state formation is described as the presence of supra-tribal government between 1CE - 1500CE. We downloaded modern socio-economic data from the World Bank data portal for total debt as a % of GDP, agricultural land %, control of corruption, rule of law, government effectiveness, political stability and absence of violence, regulatory quality, and voice and accountability. Note that the latter 6 metrics all indicate governance quality and were highly correlated, so we used only government effectiveness as a covariate as it had the highest correlation with the others. A primary motivator of European exploration was control of mineral wealth in the form of gold, silver and diamonds, as such, we counted the number of known mineral resource deposits for these three minerals per country from the Mineral Resources Data System hosted at the USGS.

We formulated the following expected paths between variables based on multiple hypotheses and findings from the published literature. From this conceptual model, we generated a structural equation model representing statistical relationships between countries’ social, geographical, natural and economic properties from historical to modern periods. Importantly for our work, we allowed a direct link between biodiversity and poverty metrics, testing the “biodiversity underpins local wealth” hypothesis. We also allowed biodiversity metrics to effect colonisation probability and state-formation probability, and biodiversity to indirectly impact poverty through the model structure, testing the “systemic bio-inequity” hypothesis. The model structure is presented in Figure 1.

We fitted this full model as shown in Figure 1. From this full model, we observed that the non-mechanistic link between latitude and each of precolonial state, colonialism and resource rent variables were not d-separated, indicating that including these links would result in a better supported model structure. Given the already oversaturated state of our model, the constrained sample sizes (ranging from 85 to 108 countries depending on the poverty metric), and lack of mechanistic process, we did not further fit these links. We also fit models where we replace our metrics of biodiversity and mineral availability with latitude to examine the results of a simpler model structure. Latitude was strongly related to the suite of factors in our SEM, with one of these being biodiversity. Yet in this model, factors are not mechanistically related to poverty and its proposed determinants (hereafter “non-mechanistic models”). As such, we also fitted a “mechanistic-only” model, which we present in the main manuscript, including biodiversity metrics and mineral resources directly and removing latitude as a predictor from all models. The logic here is that latitude is not a direct driver of any property in this model, except for biodiversity. Fitting this full model without latitude, we then performed backwards stepwise model selection iteratively removing variables until only variables with significant coefficients were retained. We calculated Fisher’s C statistic and chi-squared statistics, but caution we have a low sample size for SEMs based on the available data across all countries and instead prefer to interpret coefficient effects rather than compare amongst competing overall model structures. We calculated the direct and indirect effects of biodiversity metrics on poverty in our final models. We fitted non-mechanistic and mechanistic models for each combination of poverty metrics (n=4) and biodiversity metrics (n=3). The direct links between biodiversity and poverty metrics test the “biodiversity contributes to local wealth” hypothesis, the indirect links between biodiversity metrics and poverty through colonialism, governance and debt tests the “systemic bio-inequity hypothesis”.

We fitted structural equation models using the *piecewiseSEM* package (version 2.3.0; (Lefcheck 2016)) and calculated direct and indirect effects using 1,000 bootstrap repetitions using the *semEff* package (version 0.7.2; (Murphy 2024)).

To provide deeper contextual information to historically situate our SEMs findings, we compiled data on the history of wealth. We compiled data on the geographic location of well-established historical civilisations, empires and polities from Wikipedia with subsequent consultation of original references, focusing on those established prior to 1500s, apart from the Mughul empire (1526 BCE). Geolocation points were represented from the coordinates of reported metropolitan centres but note that many civilisations or empires spanned larger areas than represented by point coordinates. Our aim is not to be comprehensive but instead to provide a spatial overview of well-known large polities which can be interpreted in the context of modern spatial gradients in poverty and biodiversity. We compiled GDP (PPP) per capita from 1000CE to 2000CE data for West Europe, United States, India, China and Africa from Maddison (2007) “Contours of the World Economy, 1-2030 AD” (Maddison 2007).

### Comparing biodiversity, poverty and conservation

We next aimed to qualitatively categorise and quantitatively cluster our globally consistent dataset of poverty and biodiversity metrics for all 275 countries, territories and areas of geographical interest (units defined by the standard ISO 3166-1) available in the World Bank dataset (but note only 86-108 countries with complete cases of raw data were used in the SEM above). We used the dataset with 168 countries and their poverty and biodiversity metrics to impute the values for the missing metrics of all 275 ISO 3166-1 units. We performed imputation using the missForest package in R (Stekhoven & Bühlmann 2012) (verson 1.5) using default settings other than setting *maxiter* as 10,000 and *ntree* as 1000. These imputations provided excellent imputation quality with a median out-of-bag normalised root mean-squared imputation error of just 1.4%, ranging from 0-1.5%.

We first qualitatively classified countries into “high” or “low” categories for poverty and biodiversity by having the combined indices (described above) greater, or less, than 0.5. We classified conservation as “high” or “low” using a threshold of 17%, a value based on the Aichi biodiversity targets. We therefore identified countries as falling into one of 8 predefined categories being high or low for biodiversity, poverty and conservation. This subjective, but simple definition was then compared to a quantitative and data-driven analysis of clustering across all metrics of our poverty, biodiversity and conservation data. We used model-based clustering based on parameterized finite Gaussian mixture models (GMMs) using four groups, having first assessed the optimal model parameterization and number of groups using Bayesian Information Criteria (Scrucca *et al*. 2023). We also ran dimension reduction for model-based clustering to obtain the linear combinations of the original biodiversity, poverty and conservation metrics quantified by the associated eigenvalues. For each country, we therefore obtain the qualitative classification based on our subjective thresholds and the quantitative classification based on the GMMs. We then ran a chi-squares test on the country groupings between the two methods, which in all preliminary assessments, and in our final analysis was highly significant (see Results). We present this work using the World Bank’s classification of countries into 5 regions: Africa, America’s, Asia, Europe and Oceania, but acknowledge cultural and political diversity is overlooked when defining heterogeneous countries into regions.

All analyses were performed in R (version 4.3.3) for full session info and package usage see “Code Availability”.

## Results

### Biodiversity-poverty relationships at a global scale

We find no support for the negative relationship between biological diversity and poverty that would be expected under the “*biodiversity underpins local wealth*” hypothesis (Figure 2). That is not to say biodiversity does not broadly contribute to well-being, but that the emergence of a global association is not evident. Overall, we find consistently positive relationships: higher poverty and greater inequality in countries with richer biodiversity (median Spearman’s rank correlation = 0.42 with between-metric range from [0.15 to 0.58]; median R2 of simple linear regression = 0.19 [0.02 to 0.30]). This finding provides substrate to further investigate the “*systemic bio-inequity*” hypothesis. Countries’ residual deviations from the unexpected positive relationship between biodiversity and poverty indices show that much of Africa and some countries in West and SE Asia have even higher rates of poverty than predicted based on the global patterns of biodiversity, whereas national poverty is less dramatic than predicted in most of southeast Asia and south America (Figure 3).

The strength and consistency of the association that is opposite to the prediction from the “*biodiversity underpins local wealth*” hypothesis, highlights the existence of processes which decouple or modify how variation in biodiversity leads to well-being and poverty reduction. Indeed, our structural equation models revealed an indirect net effect of higher biodiversity leading to greater poverty. Biodiversity indirectly increased metrics of economic poverty, mainly through colonisation and governance strength (poverty headcount and national poverty headcount in Figure 4, Figure S2, S3). We found quantitative support for a complex set of paths from biodiversity to poverty, whereby biodiversity rich countries had higher likelihood of being colonised. Biodiversity rich countries also had lower likelihood of pre-colonisation state formation which indirectly increased colonisation rates. Colonisation increased economies dependence on the export of raw materials (higher natural resource rent) which weakened governance effectiveness. Lower governance effectiveness was associated with higher debt and higher poverty rates. Notably, mineral richness did not explain a significant portion of variation in poverty rates, which might have been expected if colonising countries were mainly exploiting mineral wealth. This overall structure in our model, and the paths, linking biodiversity to poverty was statistically supported for most metrics of biodiversity with poverty headcount and national poverty headcount (Figures S3, Table S2), but the multi-dimensional poverty index or Gini index did not show significant support in our path diagram. Replacing biodiversity metrics with latitude showed the same overall structure. Notably, a direct link between biodiversity and any poverty metric, either positive or negative, was never supported across all 12 combinations of poverty and biodiversity metrics. Structural equation modelling of our data, hence, identifies a plausible pathway to explain the net positive relationship between biodiversity and poverty.

### An unexpected inversion of biodiversity-poverty relationships given historical GDP

The uncoupling between economic poverty and biodiversity that we observe appears as a historical anomaly, with ecological and social systems being geographically more closely coupled through most of human history. Consistent with the hypothesis that “*biodiversity underpins local wealth*” is the knowledge that economies were larger and human well-being higher in major sectors of the Global South than in the Global North from prehistoric times, and throughout the pre-imperial and pre-colonial eras ((French 2021; Maddison 2007); Figure 5a). Many of the largest economies in ancient, classical and mediaeval times emerged in Africa, South and Central America and West, South and South-East Asia between 20° South and 40° North (Figure 4b; (Habib 2013; Maddison 2007)), and those in Europe emerged in Europe’s South-East and South (Minoan, Greek, Roman). Just prior to European colonialism, 20-30% of global GDP is estimated to have been generated in the Indian sub-continent, a region that now generates <5% of global GDP (Maddison 2007), and the motivation for early West European exploration of Africa was the mythical wealth of polities south of the Sahara (French 2021; Gomez 2018; Maddison 2007). That said, small and low-income societies with tight socio-ecological coupling also report high levels of life satisfaction (Galbraith *et al*. 2024). Indeed, while data is scarce and global research patterns biased, findings indicate the last 1000 years has seen a reversal, and transfer, in global proportions of wealth from Africa and India to Europe and the USA (Figure 5c). This suggests that more closely coupled socio-ecological systems have become geographically uncoupled only in the past 500 years, in the wake of European colonisation and the lasting hegemony that resulted from it, thereby inverting the direction of the relationship between biodiversity and prosperity of regions.

### Positive relationship between poverty and biodiversity in a protected world

In this context of evidence more strongly aligning with systemic bio-inequity, we next investigated whether conservation investment is distributed efficiently and justly across gradients in poverty and biodiversity. If so, we expect higher protected area coverage in regions with higher biodiversity and lower poverty (Figure 6a), with other scenarios indicating inefficiencies and injustice. These predictions assume that protected areas have costs, and that global biodiversity conservation is more efficient when protected areas are in high biodiversity regions.

We found no support that more biodiverse or less poverty-stricken regions have higher protected area coverage, despite the intersections of poverty and biodiversity we report above (Figure 6b; biodiversity: slope = 1.24, t = 1.08, p > 0.05; poverty: slope = -0.35, t = -0.24, p > 0.05). This illustrates a decoupling of conservation investment from biodiversity-poverty gradients. For example, 36% of African countries invested in protected areas with coverage greater than 17% of national land, which is within the global range across all other continents (17 - 69%). Yet, 84% of African countries had above average poverty rates, far higher than any other continent (0 - 29%), and Africa has the most countries with above average biodiversity (35%) compared to other continents (0 - 29%). In contrast, all European countries had below average poverty and biodiversity but 69% of European countries had already committed more than 17% of land to protected areas.

We find strong geographical signals in the global patterns of biodiversity, poverty and conservation investment combined: countries fall into 4 quantitatively determined groups (Figure 6c and 6d). However, none of these groupings were strongly related to protected area coverage (maximum standardised basis vector = |0.15|). The main factors contributing to these four groupings were variation in poverty rates (standardised basis vector = -0.94) and species range richness (-0.3). A second axis of variation determined grouping due to multidimensional poverty (0.54), alongside divergent gradients in species range richness (0.49) and mammal genetic diversity (-0.49).

## Discussion

### Historical-to-modern processes underpin biodiversity-poverty relationships

While acknowledging the lack of a truly causal analysis, our findings are more consistent with the “systemic bio-inequity” hypothesis, compared with the “biodiversity underpins local wealth” hypothesis, at least at a global scale. We also acknowledge these hypotheses are not mutually exclusive. Our path analysis indicates systemic bio-inequity: the benefits of biodiversity have been, and still are, systemically accumulated in privileged sectors of global society, with historical pathways helping establish this accumulation which are likely self-reinforcing to the benefit of biodiversity-poor countries of the Global North.

The latitudinal biodiversity gradient is a ubiquitous pattern in macroecology, whereby low latitudes are expected to hold more plant and animal species and populations (Pontarp *et al*. 2019). This diversity of organisms are of value to past, modern and future societies (Pironon *et al*. 2024). Unsurprisingly then, most major agricultural crops trace their origins to low latitudes, including cotton, coffee, sugarcane, tobacco, tea, rice, maize, potato, cocoa and oil palm, and many of these groups were domesticated from more than one wild species that were crossbred. Given this richness of biological resources at low latitudes, under an ecosystem service viewpoint, it is paradoxical that poverty peaks at low latitudes and prosperity peaks in the temperate North. Our work resolves this paradox by incorporating quantitative data which associates biodiversity gradients to European conquest and colonialism, from the 1500s – 1960s, and the subsequent enforcement by the former colonial powers of neoliberal free-market economic policies and unequal terms of trade on governments in the Global South. These economic policies facilitated debt creation alongside repayment structures that entrench structural poverty, political dependency and North-South inequality, giving the economically powerful Global North control over pricing of biodiversity and natural resources from the Global South (Acemoglu *et al*. 2001; Bradshaw & Huang 1991; Hickel 2018b; Robbins *et al*. 2016; Zajontz 2022).

The flow of ecosystem services from biodiversity rich countries, as proposed in the “systemic bio-inequity” hypothesis, seems a plausible mechanism explaining the positive poverty-biodiversity relationships. Our findings are entirely consistent with world-systems theory and ecologically unequal exchange whilst seeming paradoxical through the dominant lens of ecosystem service theory. In this framework, the periphery provides raw materials (biodiversity) to the core, absorbing the ecological costs while exporting the economic value. Our findings therefore add a historical and biodiversity focused perspective to the understanding that ecologically unequal exchange is driven by economic power and biophysical resources (Dorninger *et al*. 2021). As such, new models of ecologically unequal exchange could further investigate the role of biodiversity appropriation in guiding colonial interests and the resulting modern economic power structure. A network of historical-to-present day impacts were quantitatively supported in our path analysis but also aligns with previous quantitative and qualitative research. For example, our explanation aligns with historical records and current patterns in unequal exchange and trade (Hickel *et al*. 2022a; Wang *et al*. 2024; Wiedmann & Lenzen 2018). In this scenario, colonialism and imperialism violently appropriated the economic benefits of biodiversity - that in principle would help reduce poverty in regions of its origin - to benefit European nations. This process aligns with Harvey’s concept of “accumulation by dispossession”, where the commodification of nature transfers wealth from the periphery to the core (Harvey 2003), effectively fuelling Europe’s rapid economic development during the past few centuries. Positive feedbacks are then reinforced when the institutions that generate economic wealth in a neoliberal economy are environmentally extractive, rather than investing wealth back into countries’ institutions (Acemoglu *et al*. 2001), a phenomenon shown to have enduring impacts on poverty and inequality today (Aracil *et al*. 2022; Hickel *et al*. 2022a; Infante-Amate & Krausmann 2019). These mechanisms would have reduced poverty in biodiversity-poor colonising countries and thereby impoverished and underdeveloped biodiversity-rich countries in the Global South (Rodney 1974).

A specific and illustrative example of our systemic bio-inequity hypothesis comes from the evolutionary and agricultural history of cotton, which provides a deeper causal examination for the global conservation field to consider (Wendel *et al*. 2009). Wild cotton (*Gossypium* spp) is a species-rich genus of Rosid plant. Its more than 50 wild species are distributed naturally in the tropics and subtropics. Most wild cotton species are diploid, but a group of five species from Mesoamerica, the Caribbean and Pacific islands are tetraploid, leading to rapid speciation and diversification of the clade. They arose as a result of a natural hybridization event between pre-existing species around 1.5 to 2 million years ago, a case of diversity begetting more diversity, characteristic of species-rich tropical biomes (Wendel *et al*. 2009). Indigenous agriculturalists had worked with the tetraploid species that produce significantly larger and more desirable cotton bolls. They bred cultivars in Mesoamerica long before European conquests. After colonisation, local people were enslaved and their labour appropriated for mass-production of cotton for export to Europe. The genocidal enslavement of native Americans was followed by importation of enslaved labourers from Africa and expansion of the plantation system into south-eastern North America. Upon arrival in British-colonised North America, this industry sparked the development of chattel slavery which in turn fuelled the industrial revolution in Europe (Berg & Hudson 2023; French 2021; Williams 1944). Tetraploid cotton has been the most used natural fibre ever since the 1700s, amounting at times to 80% of world natural fibre production (Voora *et al*. 2023). This example serves to illustrate how the technical and economic rise of western Europe (and its settler colonies) was fuelled by the appropriation of biodiversity resources and labour from the Global South (Hornborg 2006). In addition, past and present cotton plantations impact biodiversity negatively through large scale transformation of land to monocultures across the tropics while unequal trade rules simultaneously entrench the Global North’s control over the global price for cotton (Voora *et al*. 2023). Agriculturalists continue to develop new varieties of cotton by cross-breeding with wild species that only grow in Global South countries, to incorporate desirable traits of wild species into the commercial species (e.g., pest resistance and drought tolerance; Wendel *et al*. 2009). Similar stories linking tropical evolutionary diversification, colonial exploitation and ecologically unequal exchange creating poverty in the South could be told about many other major crop plants (Paterson *et al*. 2013), contributing to Europe’s rise to dominance and the simultaneous devastation of economies and environments across the South (French 2021; Horne 2017). Many crop plants of global importance derive from highly species-rich tropical clades, and their domestication, very often involved genetic contributions multiple of species (Pironon *et al*. 2024). This points to the importance of species rich biomes, where colonialism focussed its efforts, as the source of domestic plants even when the domesticated products eventually can be grown beyond the tropics.

A first limitation to our work is that we overlook some important aspects of biodiversity, poverty, well-being and conservation by using broad scale country-level indices that are imperfect representations of natural and social systems. For instance, while the multidimensional poverty index is the most comprehensive global measure of well-being at a country level, it does not properly account for non-material aspects of wealth. Additional concepts of wealth include social capital and a sense of community, the spiritual and cultural values of nature, living a meaningful life, or a cultural sense of place that are also important constituents of well-being (Galbraith *et al*. 2024). Despite the possibility that using higher resolution data, or sub-national case-studies would modify our results and their interpretation, the global patterns we report are well-aligned with decades of qualitative assessments in interdisciplinary social sciences (Dorninger *et al*. 2021; O’Brien *et al*. 2024; Zhang *et al*. 2023). We should also be note, that despite our focus on how historical social and economic processes shape global poverty and conservation there are many modern aspects of sustainability policy that both reinforce or counter these outcomes (Kashwan 2017; Kashwan *et al*. 2021a; Koot *et al*. 2024). For example, (Kashwan *et al*. 2021a) outline modern issues with conservation practice through new conservation finance mechanisms. These initiatives often fund militarised conservation approaches or appropriate control over land from local people for conservation. As such, conservation exists in a historical structure but is not necessarily historically determined, with modern initiatives continuing to define the outcomes of systemic bio-inequity.

### A conservation paradox that urges realignment of conservation investment, poverty, and biodiversity gradients

In the context of systemic bio-inequity, we also revealed a paradox that global conservation investments, encouraged by the Convention on Biological Diversity over recent decades, are poorly aligned with the economic capacity for conservation and with the distribution of biodiversity. We found that the assumption that biodiversity underpins local wealth is inherent to the global CBD policies, however, this assumption remains unsupported here, and that protected area placements largely overlook gradients in biodiversity and poverty. Our results indicate that the share of this conservation burden is not currently well-balanced against differences in poverty and biodiversity among countries and peoples. Our results therefore support calls for Connected Conservation (Carmenta *et al*. 2023b). This framework acknowledges the tele-coupled nature of biodiversity degradation, and therefore targets wealth accumulation as a fundamental driver of biodiversity loss, which we show has benefited from systemic bio-inequity (O’Brien *et al*. 2024). Under such a framework, influential global conservation agendas under the auspices of the UN CBD and international NGOs that focus on biodiversity conservation should increasingly target the underlying, powerful and indirect drivers of global biodiversity loss such as wealth accumulation and growth centred development models (Moranta et al. 2022; Otero et al. 2020).

Our work builds on other scholars’ recognition of the important role that inequality and democratic strength play, at national and international levels, in promoting or avoiding protected areas (Kashwan 2017; West *et al*. 2006). That is, stronger democracies and greater economic equality leads to lower protected area coverage, indicating democratic processes tend to avoid separating nature from people. Our conservation paradox is problematic because the effectiveness of protected areas is highly heterogeneous, and there exists little consensus that protected areas are effective across social, economic and ecological outcomes (Brockington & Wilkie 2015; Mbaria & Ogada 2016; Zhang *et al*. 2023). However, there is broad agreement that social-ecological systems decoupling, observed here at a global scale, is a common cause of failure in nature protection (Carmenta 2023). While well designed conservation actions do benefit biological diversity in the short-term in the studies available (Langhammer *et al*. 2024; Sutherland *et al*. 2021), especially in terrestrial and marine but not generally in freshwater systems (Haase *et al*. 2025) (but note far fewer studies exist in economically poorer countries). If nature protection is advanced without reducing local poverty then conflicts are exacerbated, people dispossessed of land, and power structures favouring local elites reinforced (Brockington & Igoe 2006; Mbaria & Ogada 2016; Milne 2022; Zhang *et al*. 2023). As such, the widespread uncritical acceptance of that conservation-driven improvements to biodiversity and ecosystem-services will reduce poverty may have harmful consequences. These harms may be greater in the countries within regions having lower-than-average biodiversity and higher-than-average poverty (41% and 14% of countries in Africa and Asia respectively) and for underprivileged sectors of rural society within most countries. Global-scale conservation science and conservation planning often fail to address questions of power, politics, inclusion, or vested interests in local realities. This can lead to potentially harmful assessments of the interrelationship between people and nature (Dencik *et al*. 2019; Kashwan 2017; Pritchard *et al*. 2022; Schultz *et al*. 2022) and scientifically narrow conclusions that perpetuate unjust practices and entrench the decoupling of social and ecological systems (Rakotonarivo & Andriamihaja 2023; Wilshusen *et al*. 2002).

The separation of people from nature, most distinctly through protected areas, is a predominant mode of western conservation which decouples socio-ecological systems (Büscher & Fletcher 2019; O’Brien *et al*. 2024; Trisos *et al*. 2021). We find the extent of protected areas is largely unrelated to poverty or biodiversity which is likely symptomatic of a system that overlooks local and Indigenous people’s participation, rights, and sovereignty. Instead, this mode of conservation imposes discursive and material separation of people and their surroundings into the categories of nature, culture, environment, and society based on Eurocentric values and worldviews (Mbaria & Ogada 2016; Schultz *et al*. 2022; West *et al*. 2006; Wilshusen *et al*. 2002; Mpassi 2025). Recent shifts in the encoding of local values in conservation algorithms highlight how justice-based approaches can jointly benefit biodiversity and people (Venier-Cambron *et al*. 2024). These may eventually guide us away from global-scale conservation prioritisation schemes that lack local context (Agrawal 2021; Cobb *et al*. 2024), but in the meantime many communities in and around protected areas in the Global South continue to face dispossession and violence. We should acknowledge that conservation as a concept is far broader than protected area coverage used in our work. Our reliance on official PA coverage may overlook indigenous stewardship or community-based conservation (OECMs) that are not formally recognized by the state. However, as PA coverage remains a core metric for the 30×30 Global Biodiversity Framework target, analysing its global distribution remains critically relevant for policy critique. We use protected areas as they are widely available but note that they do represent the most extreme form of nature-people disconnections such that this metric likely exposes issues with using conservation as a poverty alleviation tool.

To provide local evidence in support of our global scale analysis (which cannot address these local complexities), we provide here specific examples of how decoupling of local socio-ecological systems through protected areas can lead to social and ecological harm. For instance, in Tanzania’s Eastern Arc Forests local communities bear significant protected area costs, while international stakeholders reap financial gains (Platts *et al*. 2023). In Namibia, extreme racialised inequalities in land ownership were inherited from the colonial regime, which still negatively impact PA effectiveness designated since the 1990s in community conservancies (Hoole & Berkes 2010). In Southeast Asia too, protected areas often overlap with territories where Indigenous and other minority groups have weak or informal tenure. States are frequently reluctant to formalize these rights, and Indigenous populations are often concerned that conservation is a means of resource appropriation by elites, as seen in Myanmar (Liljeblad 2022; Woods & Naimark 2020), Cambodia (Milne 2022) and Kenya (Mbaria & Ogada 2016). In Ethiopia, Western ideals of protected areas, assumed ecological degradation, and overpopulation in an “African Eden”, favoured the displacement of local pastoralists and the establishment of a protected UNESCO world heritage site as a solution to these perceived problems (Blanc & Morrison 2022). In Vietnam, over the last three decades, conservation and ecosystem restoration efforts have prioritised large scale, economically profitable, and quick solutions, with the unintended consequence of exacerbating biodiversity loss by converting natural forests to exotic fast-growing species that do not support long-term ecological or social integrity (McElwee 2011; Nguyen 2021; Sikor & Nguyen 2011). These examples highlight a persistent disconnect between supposed biodiversity conservation benefits from the social, political and economic realities. This disconnect is currently a key systemic driver of local conservation failures and biodiversity loss (IPBES 2019; O’Brien *et al*. 2024), and our analysis suggests that this disconnect is also systemic at the global scale.

Our findings further support calls that conservation and sustainable development policies, and decision makers across the hierarchies of global policy (Nguyen *et al*. 2025), should focus attention on the underlying causes of this paradox. Notably, the IPBES transformative change assessment is a step forwards in this direction, but uptake of global conservation policies that attempt to reduce the wealth inequalities generated by extractive capitalism, connect people to nature, and build less individualistic societies are far from realisation. First and foremost, we must recognise that long-term sustainability may be undermined by narratives that advocate for sustainable development through biodiversity conservation. These narratives risk downplaying the colonial and hegemonial structure underlying current benefit-sharing from conservation. Better recognition and understanding of these historical and current political interlinkages may help shift towards decolonising conservation science (Rakotonarivo & Andriamihaja 2023) and to shift applied conservation to forms that are globally aware but locally rooted, pluralistic and socially just (Owuor et al 2025). Conservation practise should prioritise the restoration of the coupling between social and ecological systems from local-to-global scales (Corbera *et al*. 2021). The recognition that inverted poverty-biodiversity relationships emerge from colonial, neocolonial and neoliberal economic structures urges knowledge brokers in civil society, especially those who inform government and international policy, to actively support equitable and locally-driven solutions, and recognise the limitations and historical injustice of top-down, neocolonial or Eurocentric conservation strategies (Nguyen *et al*. 2025). One specific example where scientific and conservation discourses act unjustly and ignore structural inequality is debt-for-nature swaps, whereby tweaking financial systems towards nature overlooks the injustice of Global South debt in the first instance, and instead opts for paternalistic biodiversity conservation from afar (Hassoun 2012; Losos *et al*. 2024). Indeed, the worldviews that lead to such pseudo-solutions are dominant due to existing power imbalances which promote Eurocentric ontologies and epistemologies, embedded in mainstream conservation approaches (Kashwan *et al*. 2021a). These divide people and nature through protected areas, use capitalistic market-based instruments (Büscher & Fletcher 2019; Corbera *et al*. 2021) and contain Eurocentric value-laden concepts (West *et al*. 2006; Mbembe 2022). Instead, context-specific conservation strategies adapted to governance modes, economic dynamics, cultural practices, and historical legacies will better facilitate work towards transformative solutions (Büscher & Fletcher 2019; Gurung *et al*. 2025; Kashwan *et al*. 2021b). To enable such solutions, global system-wide restoration of mechanisms supporting local social-ecological feedback, those which also benefit biodiversity, is urgent. This requires measures such as international debt-cancelling, fair commodification of biodiversity resources and establishment of fair trade to rebalance persistent ecologically unequal exchange.

## Conclusion

Our results indicate a critical framing of global conservation narratives is in order, one that acknowledges the prevalence of relations between biodiversity and poverty that are opposite to those predicted by the dominant and popular “biodiversity underpins local wealth” narrative. Further introspection is needed on how historical inequities can be placed at the centre of any globalised perspective on curtailing biodiversity loss and poverty through sustainable development and conservation, better recognising that these inequities originated from colonialism and are now entrenched in the modern global economic systems. Our work highlights that classic ecosystem service theory that “biodiversity underpins local wealth” likely predominated through much of human history, as indicated by the historic distribution of wealth and polities. However, several centuries of separation and exploitation of nature and people have led to the breakdown of social-ecological system coupling leading to systemic bio-inequity. Such an observation aligns with the theory of ecologically unequal exchange and is grounded in decades of interdisciplinary socio-ecological theory (Corsi *et al*. 2024). Countries in the Global North now leverage power to argue that conservation in the Global South, often through protected areas, would bring poverty alleviation. This occurs despite unequal global trade rules continuing to favour the rich North. A major effort is required to create a fair global trade system with the goal of restoring local-to-global social-ecological system coupling at its heart. The continued export of the idea that “biodiversity underpins local wealth” risks undermining conservation motivation in the Global South and risks implementation of unjust conservation practices, embedded in global inequalities, which is deeply unsustainable.

## Data Availability

The datasets generated during and/or analysed during the current study are available in the figshare repository and will be available after submission of the article.

## Code Availability

The code to reproduce the figures and analyses in this manuscript are available at https://github.com/cwaldock1/biodiversity-poverty which will be made a release version on acceptance.

## Author contributions statement

Author contributions following the Contributor Roles Taxonomy (CRediT):

C.W. – Conceptualisation, Data curation, Formal analysis, Investigation, Methodology, Project Administration, Software, Visualisation, Writing - Original Draft, Writing - Review and Editing

M.A.O. – Conceptualisation, Writing - Original Draft, Writing - Review and Editing

V.T.H.N. – Conceptualisation, Writing - Original Draft, Writing - Review and Editing

A.S. – Writing - Review and Editing

O.S. – Conceptualisation, Investigation, Project Administration, Writing - Original Draft, Writing - Review and Editing

## Positionality, inclusion and ethics statement

In writing a global study, we must reflect on who we are as authors and who we imagine we write for (Abimbola 2019). CW undertook his education and post-doctoral work in the Global North and, in writing on a global topic, unavoidably applied a foreign pose (from where one writes). He writes for others with a foreign gaze (for who we write) to better recognize that research on global biodiversity and poverty, and the resulting conservation “solutions”, exist in a world of power imbalances and asymmetries that shape interpretations which continue from the colonial project. OS undertook most of his education in the Global North but lived and studied in the Global South for some years in his early formation. He has continued to work with partners and students in and from the Global South throughout his career, and his family is rooted in the Global South. He writes for readers in the Global South and in the Global North, struggling with identity conflicts through much of his academic career. MAO is from Kenya, “Global South”, lived, undertook her education and developed a career in Kenya working in academia and initiating conservation projects with local communities in Kenya. MAO undertook her studies at MSc and PhD in the “Global North” through scholarships. Her exposure and experience allows her the privilege to compare the understanding she has of the inequality and power imbalances in the issues we are addressing in this manuscript between the “Global North” and “Global South”. MAO is writing from experience and lived realities for the global community of scientists, researchers, development organizations and governments both North and South with the hope that goodwill and interest prevails to address conservation and economic disparities in an integrative and inclusive way as we aim to realize “biodiversity underpins local wealth”. VTHN’s education and professional experience span diverse contexts, including mainland Southeast Asia, the USA, and Europe. This global trajectory provides a unique perspective for the paper on the ways global environmental problems like biodiversity loss manifest locally—capturing both their differences and their shared commonalities (“different but also same same”). VTHN’s work focuses on critically examining issues of power imbalances, social justice, human vulnerability, and resilience in the face of change. As an interdisciplinary social scientist, through this paper, she aims to bridge the perspectives of biological scientists with the social and political dimensions of biodiversity challenges to underscores the importance of integrating ecological-social systems for transformative change. Writing from these lived and professional experiences, VTHN seeks to contribute to the global conversation on biodiversity, poverty and conservation, emphasizing the need for inclusive, equitable solutions that address both local realities and global disparities. AS was born and raised in the Global South. She studied political science in the Global North. She has lived in the Global North for 30 years, which has shaped her critical thinking on North-South relations. During her academic career, she learned about the decision-making processes of international instruments and organisations whose legal mechanisms and implementations are characterised by a power imbalance that manifests itself in various political/economic/social issues, such as the unequal distribution of resources, the perpetuation of poverty, and the marginalisation of certain communities. Some of these issues, including the implementation of the SDGs, are highlighted in this article from an interdisciplinary perspective. AS writes mainly for the Global South and is committed to bringing these specific global injustices to the attention of those most affected by this systemic inequality.

## Acknowledgements

We express our gratitude to Mordecai Ogada, Carlos Melian, Mazin Qumsiyeh, Dario Josi, Bernhard Wegscheider, Marcel Hasler, Jacob Bedford Madhav Thakur, who provided thoughtful feedback on earlier versions of this manuscript and members of the Aquatic Ecology and Evolution group who have given valuable feedback in frequent discussions of this work.

## Notes

### Competing Interest Statement

The authors have declared no competing interest.

### Summary of Updates

Update based on peer review process. We made only text additions to the introduction, methods and discussion. We restructured the manuscript according to the journals formatting requirements.

