## Supporting information for "Systemic bio-inequity links poverty to biodiversity and induces a conservation paradox"

**Supporting figures and tables**

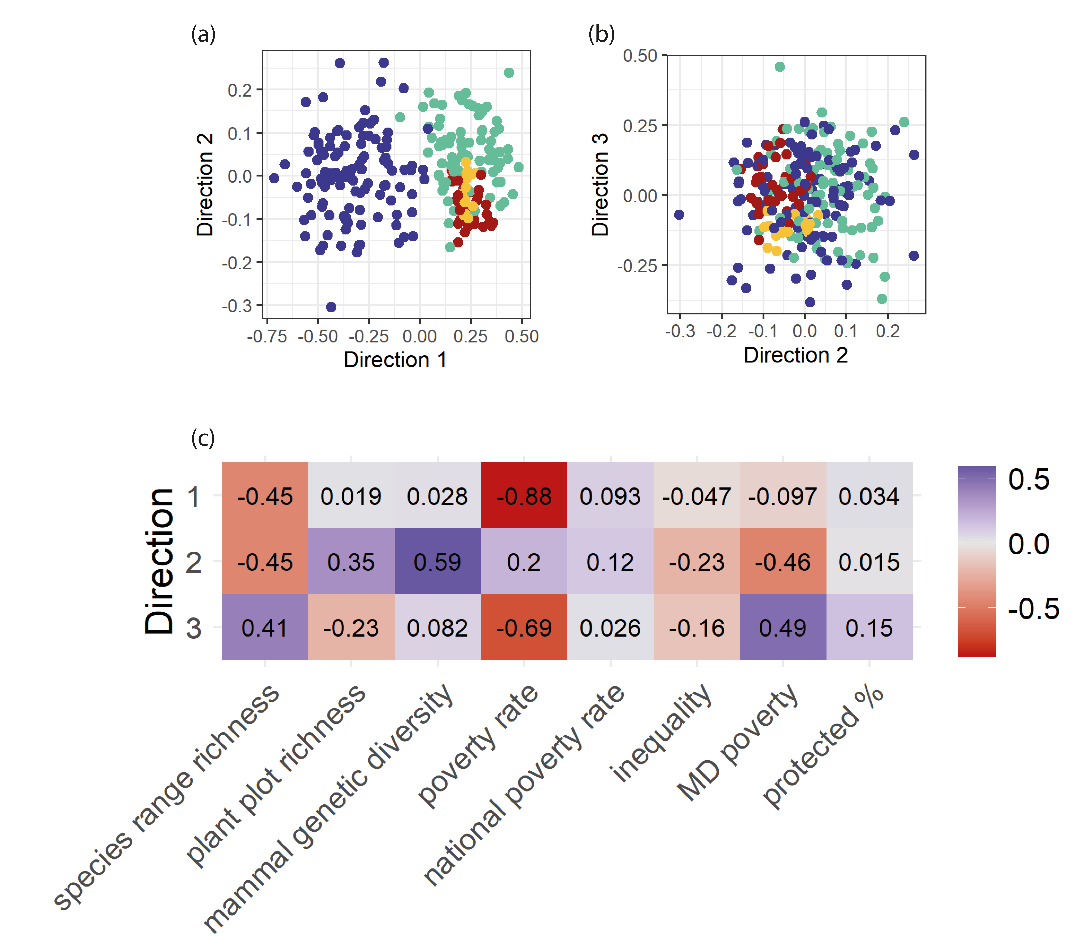

**Figure S1. Visualization of the underlying structure of the GMM subspace in three directions**. Direction 1 explains 90% of the variation, direction 2 explains 8.4% and direction 3 explains 1.5%. Panel a and b show each country plotted in this reduced subspace which was determined by the 8 metrics of poverty, biodiversity and conservation. The standardised basis of the estimated subspace reduction is represented in panel c, indicating which variables contribute to each subspace direction and the magnitude of contributions of these vectors, standardised to unit standard deviation.

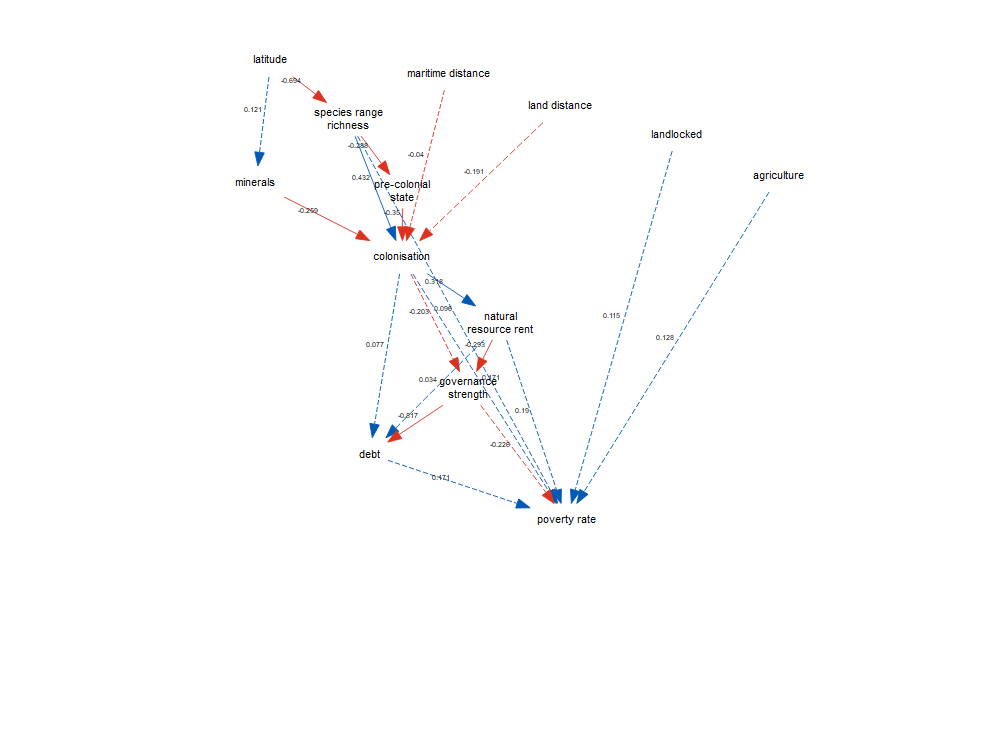

**Figure S2. Example path diagram of fitted species range richness, poverty and socio-economic relationships in a global socio-economic structure for poverty rate.** Blue links indicate positive relationships, red links negative relationships. Standardized effect sizes are indicated next to paths. Solid lines indicate significant effects of paths (coefficients). The focal links here are those between biodiversity (species range richness) and poverty (poverty rate). For all links across all poverty (n=4) and biodiversity (n=3) metrics, indicating both indirect and direct, see Figure 3 in the main manuscript and Figure S3.

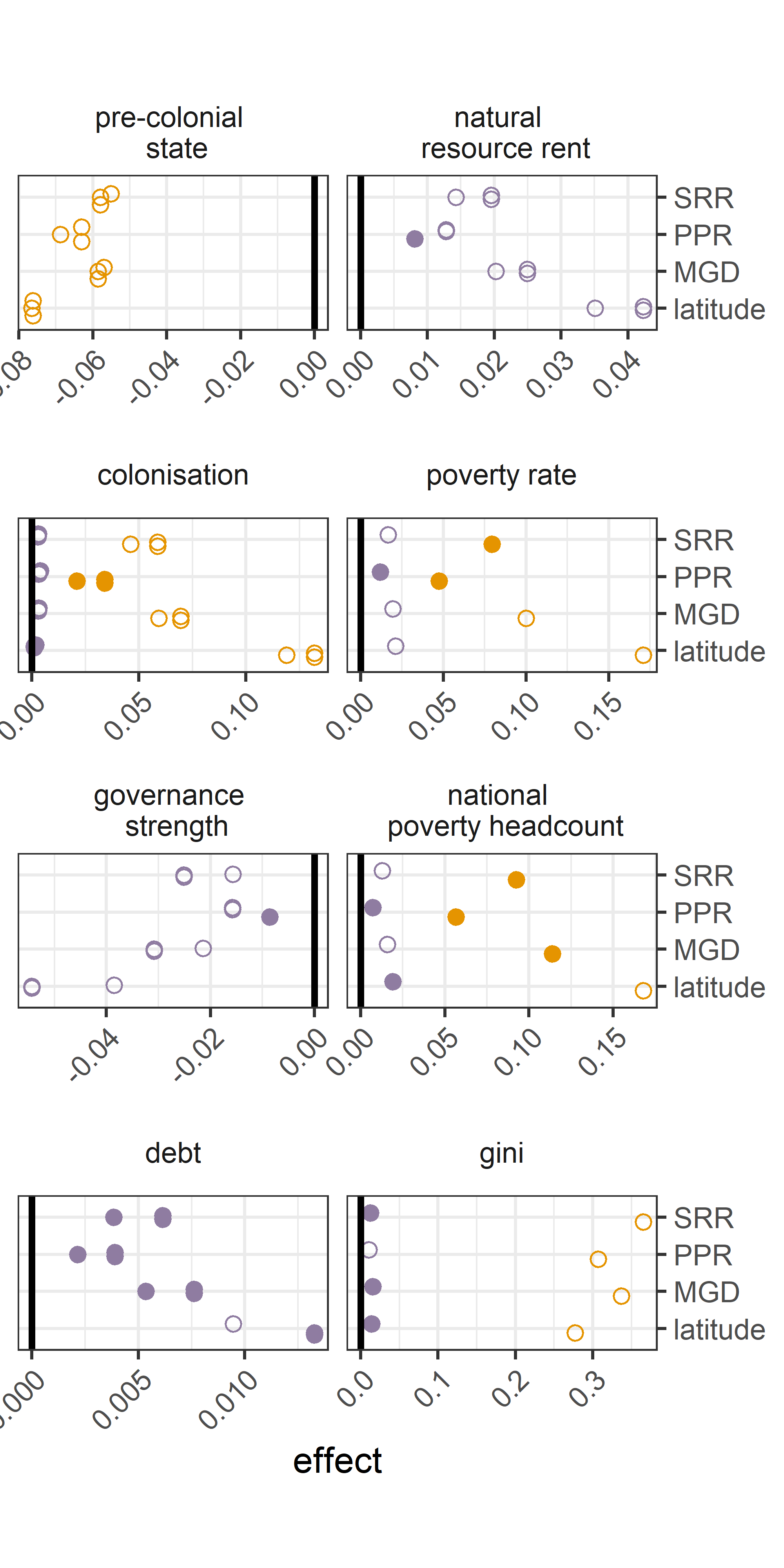

**Figure S3. Magnitude of standardized effects of biodiversity metrics on poverty responses, for both direct and indirect effects.** Effects are estimated through bootstrap resampling of the full SEM, with no backwards selection. Filled points indicate significant effects with confidence intervals from 1,000 bootstraps not overlapping 0, and empty points indicate non-significant effects. Orange points indicate direct effects and orange points indicate indirect effects. We show our model for latitude as an indicator of biodiversity gradients more broadly. *Biodiversity variables are indicated as SRR = species range richness, PPR = plant plot richness, MGD = mammal genetic diversity.*

**Table S1. Summary of data used in the manuscript and short descriptions and source links.**

|  | **Name** | **Description** | **Source** |
| --- | --- | --- | --- |
| **Biodiversity** | **Species range richness** | Terrestrial species richness for amphibians, birds, mammals and reptiles by summing overlapping range polygons to a global harmonised raster layer. | IUCN Red List Data 2022-2. <https://www.iucnredlist.org/resources/spatial-data-download> |
|  | **Plant plot richness** | Spatial model of plant local scale species richness based on >400,000 vegetation plots in a small-scale delimited area. | Sabatini, F. M. *et al.* Global patterns of vascular plant alpha diversity. *Nat. Commun.* **13**, 4683 (2022). |
|  | **Mammal genetic diversity** | Spatial model of terrestrial mammal genetic diversity of co1 and cytb mitochondrial genes comparing the mean number of nucleotide differences per site within a species for >2,000 species and >54,000 sequences. | Theodoridis, S. *et al.* Evolutionary history and past climate change shape the distribution of genetic diversity in terrestrial mammals. *Nat. Commun.* **11**, 2557 (2020). |
| **Poverty** | **Poverty headcount** | % of the country living below the international poverty line of $2.15/day based on 2017 Purchasing Power Parity | <https://data.worldbank.org/indicator/SI.POV.DDAY> |
|  | **National poverty headcount** | % of the country living below country-specific national poverty lines provided by household surveys by each country's government sources or World Bank staff which reflect cost of living differences between countries | <https://data.worldbank.org/indicator/SI.POV.NAHC> |
|  | **Multi-dimensional poverty index** | A metric of poverty that equally weighs monetary poverty, school enrolment rates, primary education completion, access to clean drinking water, access to sanitation and household electricity supply. The combined poverty index, individuals are considered poor if in the lowest ⅓ of the combined weighted index, and the country metric is expressed as the % of individuals indicated as poor. | <https://www.worldbank.org/en/topic/poverty/brief/multidimensional-poverty-measure> |
|  | **Gini index** | Gini index measures the extent to which the distribution of income (or, in some cases, consumption expenditure) among individuals or households within an economy deviates from a perfectly equal distribution. A Lorenz curve plots the cumulative percentages of total income received against the cumulative number of recipients, starting with the poorest individual or household. The Gini index measures the area between the Lorenz curve and a hypothetical line of absolute equality, expressed as a percentage of the maximum area under the line. Thus a Gini index of 0 represents perfect equality, while an index of 100 implies perfect inequality. | <https://data.worldbank.org/indicator/SI.POV.GINI> |
| **Conservation** | **Terrestrial protected area coverage** | Terrestrial protected areas are totally or partially protected areas of at least 1,000 hectares that are designated by national authorities as scientific reserves with limited public access, national parks, natural monuments, nature reserves or wildlife sanctuaries, protected landscapes, and areas managed mainly for sustainable use. Marine areas, unclassified areas, littoral (intertidal) areas, and sites protected under local or provincial law are excluded. | <https://data.worldbank.org/indicator/ER.LND.PTLD.ZS>. World Database on Protected Areas ( WDPA ) where the compilation and management is carried out by United Nations Environment World Conservation Monitoring Centre ( UNEP-WCMC ) in collaboration with governments, non-governmental organizations, academia and industry. The data is available online through the Protected Planet website ([protectedplanet.net](https://www.protectedplanet.net/)). |
| **Historical socio-economic metrics** | **Colonisation** | An indicator of whether a country was colonised by England, France, Germany, Italy, Netherlands, Portugal or Spain between 1462 and 1945. Colonisation is defined as > 20% land area under control of these countries. | Ertan, A., Fiszbein, M. & Putterman, L. Who was colonized and when? A cross-country analysis of determinants. *European Economic Review* **83**, 165–184 (2016). |
|  | **State history** | Index of the presence of supra-tribal government covering years 1CE - 1500CE. The index integrates the following information: i) the existence of a state, ii) the state being domestically based vs. imposed by an external power, and iii) the territorial extent and unity. The index is integrated and discounted such that states present nearer 1500CE are weighted more heavily. | Ertan et al. (2016)  Borcan, Oana, Olsson, Ola, Putterman, Louis, 2014. State History and Economic Development: Evidence from Six Millennia. Brown University Department of Economics, Working Paper 2014–18. |
|  | **Mineral wealth** | Sum of the number of gold, silver and diamond deposits that are known within a countries borders. We assumed a major motivator of colonising countries was to obtain access to these mineral resources perceived as valuable in Europe. We did not consider the size or quality of such deposits. We extracted all mineral resource deposits from the Mineral Resource Data System, obtained all deposits containing gold (AU), silver (AG) and diamonds (GEM_D) and summed these per country. | <https://mrdata.usgs.gov/mrds/> |
| **Geography** | **Maritime navigational distance** | Ertan et al. (2016) considered the distance between Camaret-sur-mer and the nearest port of historical significance (usually the main port) in each country. They did not consider routes going through the Suez or Panama canals. | Ertan et al. (2016) |
|  | **Land distance** | The distance that colonising countries had to travel by ground transportation once landing in an oceanic port. For landlocked countries, it indicates the distance from the country's historically most important city (usually but not always the current capital) to the closest oceanic port. For El Salvador, Ecuador, Peru, and Chile, land distance is the distance between Panama City and Balboa—the Atlantic and Pacific ports that are now joined by the Panama Canal. | Ertan et al. (2016) |
|  | **Landlocked** | Indicates if a country has direct access to the ocean. | Ertan et al. (2016) |
| **Modern socio-economic metrics** | **Government effectiveness** | Government Effectiveness captures perceptions of the quality of public services, the quality of the civil service and the degree of its independence from political pressures, the quality of policy formulation and implementation, and the credibility of the government's commitment to such policies. Estimate gives the country's score on the aggregate indicator, in units of a standard normal distribution, i.e. ranging from approximately -2.5 to 2.5. | "The Worldwide Governance Indicators: Methodology and Analytical Issues". World Bank Policy Research Working Paper No. 5430 (http://papers.ssrn.com/sol3/papers.cfm?abstract_id=1682130). |
|  | **Agricultural land** | Agricultural land refers to the share of land area that is arable, under permanent crops, and under permanent pastures. Arable land includes land defined by the FAO as land under temporary crops (double-cropped areas are counted once), temporary meadows for mowing or for pasture, land under market or kitchen gardens, and land temporarily fallow. | <https://data.worldbank.org/indicator/AG.LND.AGRI.ZS> |
|  | **Total debt service (% of exports of goods, services and primary income)** | Total debt service to exports of goods, services and primary income. Total debt service is the sum of principal repayments and interest actually paid in currency, goods, or services on long-term debt, interest paid on short-term debt, and repayments (repurchases and charges) to the IMF. This value is converted to a binary variable as whether a country holds IMF debt due to the complex structure of each debt arrangement and the national economic consequences depending on additional factors such as a countries growth rate. | <https://data.worldbank.org/indicator/DT.TDS.DECT.EX.ZS> |
|  | **Total natural resource rents (% GDP)** | Total natural resources rents are the sum of oil rents, natural gas rents, coal rents (hard and soft), mineral rents, and forest rents. The estimates of natural resources rents are calculated as the difference between the price of a commodity and the average cost of producing it. This is done by estimating the price of units of specific commodities and subtracting estimates of average unit costs of extraction or harvesting costs. These unit rents are then multiplied by the physical quantities countries extract or harvest to determine the rents for each commodity as a share of gross domestic product (GDP). | <https://data.worldbank.org/indicator/NY.GDP.TOTL.RT.ZS> |

**Table S2. Statistical summary of structural equation models terms for backward selected models.**

| **Biodiversity** | **Poverty** | **Response** | **Predictor** | **Estimate** | **S.E.** | **d.f.** | **p-value** |  |
| --- | --- | --- | --- | --- | --- | --- | --- | --- |
| **Mammal genetic diversity** | **Gini inequality** | Pre-history state | Mammal genetic diversity | -134.221 | 55.007 | 106 | 0.015 | * |
|  |  | Colonisation | Pre-history state | -3.056 | 0.869 | 103 | 0.000 | *** |
|  |  | Colonisation | Land distance | -2.131 | 1.062 | 103 | 0.045 | * |
|  |  | Colonisation | Mineral resources | -0.684 | 0.280 | 103 | 0.015 | * |
|  |  | Colonisation | Mammal genetic diversity | 318.201 | 83.456 | 103 | 0.000 | *** |
|  |  | Resource rent | Colonisation | 1.391 | 0.370 | 106 | 0.000 | *** |
|  |  | Governance effectiveness | Resource rent | -0.386 | 0.121 | 106 | 0.001 | ** |
|  |  | Debt | Governance effectiveness | -13.743 | 2.981 | 106 | 0.000 | *** |
|  | **MD poverty** | Colonisation | Maritime distance | -0.217 | 0.089 | 81 | 0.015 | * |
|  |  | Colonisation | Land distance | -3.086 | 1.162 | 81 | 0.008 | ** |
|  |  | Colonisation | Mammal genetic diversity | 197.091 | 89.099 | 81 | 0.027 | * |
|  |  | MD_poverty | Governance effectiveness | -5.141 | 1.724 | 83 | 0.003 | ** |
|  | **National poverty headcount** | Pre-history state | Mammal genetic diversity | -128.614 | 55.378 | 100 | 0.020 | * |
|  |  | Colonisation | Pre-history state | -3.064 | 0.907 | 98 | 0.001 | *** |
|  |  | Colonisation | Land distance | -2.244 | 0.982 | 98 | 0.022 | * |
|  |  | Colonisation | Mammal genetic diversity | 269.663 | 82.607 | 98 | 0.001 | ** |
|  |  | Resource rent | Colonisation | 1.308 | 0.383 | 100 | 0.001 | *** |
|  |  | Governance effectiveness | Resource rent | -0.354 | 0.125 | 100 | 0.005 | ** |
|  |  | Debt | Governance effectiveness | -12.763 | 3.006 | 100 | 0.000 | *** |
|  |  | National poverty headcount | Governance effectiveness | -1.965 | 0.970 | 100 | 0.043 | * |
|  | **Poverty headcount** | Pre-history state | Mammal genetic diversity | -134.221 | 55.007 | 106 | 0.015 | * |
|  |  | Colonisation | Pre-history state | -3.056 | 0.869 | 103 | 0.000 | *** |
|  |  | Colonisation | Land distance | -2.131 | 1.062 | 103 | 0.045 | * |
|  |  | Colonisation | Mineral resources | -0.684 | 0.280 | 103 | 0.015 | * |
|  |  | Colonisation | Mammal genetic diversity | 318.201 | 83.456 | 103 | 0.000 | *** |
|  |  | Resource rent | Colonisation | 1.391 | 0.370 | 106 | 0.000 | *** |
|  |  | Governance effectiveness | Resource rent | -0.386 | 0.121 | 106 | 0.001 | ** |
|  |  | Debt | Governance effectiveness | -13.743 | 2.981 | 106 | 0.000 | *** |
|  |  | Poverty headcount | Governance effectiveness | -4.148 | 1.400 | 106 | 0.003 | ** |
| **Plant alpha diversity** | **Gini inequality** | Pre-history state | Plant alpha diversity | -0.052 | 0.021 | 106 | 0.014 | * |
|  |  | Colonisation | Pre-history state | -2.704 | 0.793 | 104 | 0.001 | *** |
|  |  | Colonisation | Mineral resources | -0.647 | 0.258 | 104 | 0.012 | * |
|  |  | Colonisation | Plant alpha diversity | 0.069 | 0.026 | 104 | 0.008 | ** |
|  |  | Resource rent | Colonisation | 1.391 | 0.370 | 106 | 0.000 | *** |
|  |  | Governance effectiveness | Resource rent | -0.386 | 0.121 | 106 | 0.001 | ** |
|  |  | Debt | Governance effectiveness | -13.743 | 2.981 | 106 | 0.000 | *** |
|  | **MD poverty** | Pre-history state | Plant alpha diversity | -0.056 | 0.023 | 83 | 0.015 | * |
|  |  | Colonisation | Maritime distance | -0.307 | 0.092 | 82 | 0.001 | *** |
|  |  | Colonisation | Plant alpha diversity | 0.101 | 0.033 | 82 | 0.002 | ** |
|  |  | MD_poverty | Governance effectiveness | -5.141 | 1.724 | 83 | 0.003 | ** |
|  | **National poverty headcount** | Pre-history state | Plant alpha diversity | -0.055 | 0.022 | 100 | 0.011 | * |
|  |  | Colonisation | Pre-history state | -3.611 | 0.873 | 99 | 0.000 | *** |
|  |  | Colonisation | Land distance | -2.248 | 0.890 | 99 | 0.012 | * |
|  |  | Resource rent | Colonisation | 1.308 | 0.383 | 100 | 0.001 | *** |
|  |  | Governance effectiveness | Resource rent | -0.354 | 0.125 | 100 | 0.005 | ** |
|  |  | Debt | Governance effectiveness | -12.763 | 3.006 | 100 | 0.000 | *** |
|  |  | National poverty headcount | Governance effectiveness | -1.965 | 0.970 | 100 | 0.043 | * |
|  | **Poverty headcount** | Pre-history state | Plant alpha diversity | -0.052 | 0.021 | 106 | 0.014 | * |
|  |  | Colonisation | Pre-history state | -2.704 | 0.793 | 104 | 0.001 | *** |
|  |  | Colonisation | Mineral resources | -0.647 | 0.258 | 104 | 0.012 | * |
|  |  | Colonisation | Plant alpha diversity | 0.069 | 0.026 | 104 | 0.008 | ** |
|  |  | Resource rent | Colonisation | 1.391 | 0.370 | 106 | 0.000 | *** |
|  |  | Governance effectiveness | Resource rent | -0.386 | 0.121 | 106 | 0.001 | ** |
|  |  | Debt | Governance effectiveness | -13.743 | 2.981 | 106 | 0.000 | *** |
|  |  | Poverty headcount | Governance effectiveness | -4.148 | 1.400 | 106 | 0.003 | ** |
| **Species range richness** | **Gini inequality** | Pre-history state | Species range richness | -0.003 | 0.001 | 104 | 0.016 | * |
|  |  | Colonisation | Pre-history state | -2.592 | 0.823 | 102 | 0.002 | ** |
|  |  | Colonisation | Mineral resources | -0.579 | 0.264 | 102 | 0.028 | * |
|  |  | Colonisation | Species range richness | 0.005 | 0.001 | 102 | 0.000 | *** |
|  |  | Resource rent | Colonisation | 1.290 | 0.377 | 104 | 0.001 | *** |
|  |  | Governance effectiveness | Resource rent | -0.374 | 0.123 | 104 | 0.002 | ** |
|  |  | Debt | Governance effectiveness | -13.289 | 2.943 | 104 | 0.000 | *** |
|  | **MD poverty** | Pre-history state | Species range richness | -0.003 | 0.001 | 83 | 0.034 | * |
|  |  | Colonisation | Maritime distance | -0.240 | 0.093 | 81 | 0.010 | ** |
|  |  | Colonisation | Land distance | -2.743 | 1.164 | 81 | 0.019 | * |
|  |  | Colonisation | Species range richness | 0.005 | 0.002 | 81 | 0.010 | ** |
|  |  | MD_poverty | Governance effectiveness | -5.141 | 1.724 | 83 | 0.003 | ** |
|  | **National poverty headcount** | Pre-history state | Species range richness | -0.002 | 0.001 | 98 | 0.024 | * |
|  |  | Colonisation | Pre-history state | -2.878 | 0.908 | 96 | 0.002 | ** |
|  |  | Colonisation | Land distance | -2.062 | 0.944 | 96 | 0.029 | * |
|  |  | Colonisation | Species range richness | 0.004 | 0.002 | 96 | 0.007 | ** |
|  |  | Resource rent | Colonisation | 1.179 | 0.392 | 98 | 0.003 | ** |
|  |  | Governance effectiveness | Resource rent | -0.337 | 0.127 | 98 | 0.008 | ** |
|  |  | Debt | Governance effectiveness | -12.150 | 2.965 | 98 | 0.000 | *** |
|  |  | National poverty headcount | Governance effectiveness | -1.980 | 1.000 | 98 | 0.048 | * |
|  | **Poverty headcount** | Pre-history state | Species range richness | -0.003 | 0.001 | 104 | 0.016 | * |
|  |  | Colonisation | Pre-history state | -2.592 | 0.823 | 102 | 0.002 | ** |
|  |  | Colonisation | Mineral resources | -0.579 | 0.264 | 102 | 0.028 | * |
|  |  | Colonisation | Species range richness | 0.005 | 0.001 | 102 | 0.000 | *** |
|  |  | Resource rent | Colonisation | 1.290 | 0.377 | 104 | 0.001 | *** |
|  |  | Governance effectiveness | Resource rent | -0.374 | 0.123 | 104 | 0.002 | ** |
|  |  | Debt | Governance effectiveness | -13.289 | 2.943 | 104 | 0.000 | *** |
|  |  | Poverty headcount | Governance effectiveness | -4.090 | 1.413 | 104 | 0.004 | ** |
| **Latitude** | **Gini inequality** | Pre-history state | Latitude | 0.050 | 0.016 | 104 | 0.002 | ** |
|  |  | Colonisation | Latitude | -0.177 | 0.033 | 104 | 0.000 | *** |
|  |  | Resource rent | Colonisation | 1.290 | 0.377 | 104 | 0.001 | *** |
|  |  | Governance effectiveness | Resource rent | -0.374 | 0.123 | 104 | 0.002 | ** |
|  |  | Debt | Governance effectiveness | -13.289 | 2.943 | 104 | 0.000 | *** |
|  | **MD poverty** | Pre-history state | Latitude | 0.049 | 0.020 | 83 | 0.016 | * |
|  |  | Colonisation | Maritime distance | -0.229 | 0.099 | 82 | 0.021 | * |
|  |  | Colonisation | Latitude | -0.159 | 0.040 | 82 | 0.000 | *** |
|  |  | MD_poverty | Governance effectiveness | -5.141 | 1.724 | 83 | 0.003 | ** |
|  | **National poverty headcount** | Pre-history state | Latitude | 0.050 | 0.016 | 98 | 0.002 | ** |
|  |  | Colonisation | Latitude | -0.165 | 0.032 | 98 | 0.000 | *** |
|  |  | Resource rent | Colonisation | 1.179 | 0.392 | 98 | 0.003 | ** |
|  |  | Governance effectiveness | Resource rent | -0.337 | 0.127 | 98 | 0.008 | ** |
|  |  | Debt | Governance effectiveness | -12.150 | 2.965 | 98 | 0.000 | *** |
|  |  | National poverty headcount | Governance effectiveness | -1.980 | 1.000 | 98 | 0.048 | * |
|  | **Poverty headcount** | Pre-history state | Latitude | 0.050 | 0.016 | 104 | 0.002 | ** |
|  |  | Colonisation | Latitude | -0.177 | 0.033 | 104 | 0.000 | *** |
|  |  | Resource rent | Colonisation | 1.290 | 0.377 | 104 | 0.001 | *** |
|  |  | Governance effectiveness | Resource rent | -0.374 | 0.123 | 104 | 0.002 | ** |
|  |  | Debt | Governance effectiveness | -13.289 | 2.943 | 104 | 0.000 | *** |
|  |  | Poverty headcount | Governance effectiveness | -4.090 | 1.413 | 104 | 0.004 | ** |
